# The AhR is a Critical Regulator of the Pulmonary Response to Cannabis Smoke

**DOI:** 10.1101/2025.06.24.660596

**Authors:** Emily T. Wilson, Roham Gorgani, Nicole S. Heimbach, Alexandra Bartolomucci, Thupten Tsering, Julia V. Burnier, David H. Eidelman, Carolyn J. Baglole

## Abstract

Cannabis use is prevalent worldwide, with smoking being the most common method of consumption. When smoking cannabis, users are exposed to both harmful combustion products as well as cannabinoids such as tetrahydrocannabinol (THC) and cannabidiol (CBD). THC and CBD have purported anti-inflammatory effects through activation of cannabinoid receptors; however, the minimal expression of these receptors in lung tissue suggests that respiratory effects of cannabis may be mediated through alternative pathways. One potential mediator of these effects is the aryl hydrocarbon receptor (AhR), a transcription factor involved in xenobiotic metabolism. Notably, the AhR is activated by both combustion products and cannabinoids. This receptor is also known to dampen lung inflammation induced by tobacco smoke or air pollution. Therefore, we hypothesized that AhR activation would reduce susceptibility to the harmful effects of inhaled cannabis smoke. To investigate this hypothesis, *Ahr^+/-^* and *Ahr^-/-^* mice were exposed to air or cannabis smoke using a controlled puff regimen over a three-day period. In the first study to characterize the effects of cannabis smoke on lung tissue and the pulmonary secretome, including extracellular vesicles and secreted proteins, we found that acute exposure induced neutrophilia, vascular leakage, and activation of tissue remodeling pathways, all of which were regulated by AhR. These findings highlight not only the detrimental effects of cannabis smoke on lung health but also the pivotal role of the AhR as a key regulator of the pulmonary response to cannabis smoke exposure.

## Introduction

Cannabis is the third most commonly used controlled substance worldwide, with an estimated 192 million users globally (1). Growing acceptance of cannabis has led to legalization in an increasing number of countries including Canada and Uruguay (2, 3). This shift correlates with rising consumption rates (4), emphasizing the need for a deeper understanding of the health effects of cannabis. While cannabis can be consumed through various methods, smoking the dried flower, typically in the form of joints, remains the predominant mode of use (5). Cannabis smoke contains hundreds of compounds, including polycyclic aromatic hydrocarbons (PAHs) and other carcinogenic, mutagenic, and teratogenic combustion products (6), which are also present in tobacco smoke. While the link between tobacco smoking and diseases such as chronic obstructive pulmonary disease (COPD) and lung cancer is well established (7), the impacts of smoking cannabis on lung health remain unclear (6).

Although tobacco and cannabis smoke contain similar combustion products, cannabis smoke uniquely contains phytochemicals called cannabinoids (6). The most prevalent cannabinoids in cannabis smoke are tetrahydrocannabinol (THC) and cannabidiol (CBD) (8). THC is primarily responsible for the psychoactive effects of cannabis, while both THC and CBD contribute to its physiological effects, including analgesia and modulation of inflammation (9, 10). Cannabinoids exert many of their effects by binding to cannabinoid receptors 1 and 2 (CB1, CB2), both of which are G protein-coupled receptors (GPCRs) (11). However, CB1 and CB2 have minimal expression on structural cells of the lung, raising the possibility that cannabinoids could act through other receptors within the respiratory system. One such receptor is the aryl hydrocarbon receptor (AhR), which is highly expressed in barrier organs, including the lungs, and can be activated by both THC and CBD (12, 13). The AhR is a ligand-activated transcription factor that is best-known for its role in xenobiotic metabolism (14). The receptor is activated by a myriad of exogenous ligands including dioxins and combustion products as well as endogenous ligands that are tryptophan metabolites or microflora-derived compounds (15). Upon ligand binding in the cytoplasm (16), the AhR translocates to the nucleus, dimerizes with the aryl hydrocarbon nuclear translocator (ARNT), and binds to xenobiotic response elements in DNA, initiating transcription of its target genes (17). These include genes encoding xenobiotic-metabolizing enzymes, such as cytochrome P450 (CYP) enzymes CYP1A1 and CYP1B1 (18). Beyond its role in xenobiotic metabolism, the AhR also regulates genes of proteins involved in the antioxidant response such as NAD(P)H Quinone Dehydrogenase 1 (NQO1) (19, 20). More broadly, the AhR protects the lungs from damage caused by various harmful stimuli, such as tobacco smoke and air pollution by suppressing oxidative stress (20–22) and inflammation (23–28).

Since the AhR protects the lungs from smoke-induced inflammation and consequent damage (23–28) and is activated by combustion products as well as THC and CBD (12, 13), we hypothesized that this receptor plays an important role in the pulmonary response to cannabis smoke. To test this, we analyzed multiple compartments of the pulmonary system to assess the interplay between AhR and the potential adverse effects caused by cannabis smoke. Lung tissue is the structural foundation of respiratory health, such that even subtle cellular changes can directly impact lung function. Meanwhile, intercellular communication through the secretome is now recognized as an important indicator of biological function (29). We therefore collectively investigated the pulmonary secretome, which consists of extracellular vesicles (EVs) and the bronchoalveolar lavage fluid (BALF) in addition to the lung tissue. EVs are lipid-bound particles released from cells and carry molecular cargo that can modulate immune responses and tissue repair processes (30). BALF, on the other hand, is found within the lumen of the respiratory system and provides insight into the soluble factors present within the airways, offering a snapshot of the immediate response to environmental exposures (31). Our findings across these compartments reveal that cannabis smoke exposure activates AhR, which orchestrates a protective response by controlling inflammation, tissue remodeling, and metabolism.

## Materials and Methods

### Animals

*Ahr*-knockout (*Ahr*^-/-^; B6.129-*Ahr*^tm1^/J) C57BL/6 mice were obtained from Jackson Laboratory (Bar Harbor, ME) and bred in-house. A breeding scheme of *Ahr^+/-^* to *Ahr^+/-^* was used to produce offspring including *Ahr*^+/+^, *Ahr^+/-^*, and *Ahr^-/-^* mice. Male and female mice aged 8 to 12 weeks were used in these experiments. All procedures were approved by the McGill University Animal Care Committee and carried out in accordance with the Canadian Council on Animal Care guidelines.

### Cannabis products

Indica-THC dominant cannabis containing 20-26% THC and 0–0.1% CBD was purchased from the *Société québécoise du cannabis* (SQDC) (#688083002328; Quebec, Canada). Cannabis joints were hand-rolled by grinding the dried cannabis flower with a plastic grinder and packing the product into classic 1 1/4 size rolling paper (RAW®). Each cannabis joint contained 0.5 ± 0.05 g of cannabis.

### Animal exposures

*Ahr^+/-^* and *Ahr^-/-^* mice were allocated to one of two groups: air or cannabis smoke. Air-exposed mice did not receive any exposure in the system. Exposures were performed using the SCIREQ® inExpose™ system equipped with a single cigarette chamber extension. Exposure parameters and the puff profile were programmed using the flexiWare software. Puff regimens consisted of three puffs per minute with a puff volume of 35 ml. Mice were exposed to cannabis smoke twice per day for three days, with each exposure consisting of two joints. There were four hours between exposure sessions. On day three, mice were sacrificed immediately after the second exposure.

### BALF collection

BALF was collected by lavaging the lungs twice with 0.5 ml of cold PBS. The BALF was centrifuged at 3000 *g* for five minutes to pellet cells, and the supernatant was kept for multiplex analysis and EV enrichment.

### Differential cell counts

BALF cells were resuspended in PBS, counted, mounted onto slides using a CytoSpin (Thermo Scientific; Waltham, MA, USA), and stained using Three Step Stain (Thermo Scientific; Waltham, MA, USA). Differential cell counts (at least 300 cells/sample) were performed after CytoSpin slide preparation and staining.

### Isolation and characterization of EVs

Cell-free BALF was transferred to 1 ml polycarbonate tubes (Beckman Coulter; Montreal, QC) and ultracentrifuged using TLA 120.2 in a Beckman Coulter Optima MAX-XP at 110,000 *g* for 60 minutes at 4°C. Remaining cell-free EV-free BALF was removed for proteomic analysis. The EV pellet was washed with PBS and ultracentrifuged again at 110,000 *g* for 60 minutes at 4°C before protein isolation. EVs underwent nanoparticle tracking analysis (NTA) and were imaged using Transmission Electron Microscopy (TEM). For NTA, 10 μl of EVs were diluted into 490 μl of PBS and analyzed on a Nanosight NS500 system (Nanosight Ltd.; Amesbury, UK), with concentration and size distribution assessed using the NTA1.3 software (Malvern Panalytical; Montreal, QC). For TEM, EVs were fixed with 2.5% glutaraldehyde fixation solution (250 ml EMS, 50 ml 25% glutaraldehyde, 250 ml water). TEM copper grids were placed on a drop (20 μl of fixed EVs) facing down and left to settle for 20 minutes. TEM grids were washed with Dulbecco’s PBS and stained with 2% uranyl acetate for 3 minutes and air-dried overnight. EV images were taken using an FEI Tecnai™ G2 Spirit BioTwin 120 kV Cryo-TEM.

### Proteomic analysis

Lung tissue, EVs, and BALF protein samples were analyzed using LC–MS/MS. Each sample was loaded onto a single stacking gel band to remove lipids, detergents, and salts. The single gel band containing all proteins was reduced with DTT, alkylated with iodoacetic acid, and digested with trypsin. Two µg of extracted peptides were re-solubilized in 0.1% aqueous formic acid and loaded onto a Thermo Acclaim Pepmap (Thermo, 75 µm ID × 2 cm C18 3 µm beads) precolumn and then onto an Acclaim Pepmap Easyspray (Thermo, 75 µm × 15 cm with 2 µm C18 beads) analytical column for separation using a Dionex Ultimate 3000 uHPLC at 250 nl/min with a gradient of 2%– 35% organic (0.1% formic acid in acetonitrile) over 3 hours. Peptides were analyzed using a Thermo Orbitrap Fusion mass spectrometer operating at 120,000 resolution (FWHM in MS1), with HCD sequencing (15,000 resolution) at top speed for all peptides with a charge of 2+ or greater. The raw data were converted into Mascot generic format (*.mgf format) for searching using the Mascot 2.6.2 search engine (Matrix Science) against mouse protein sequences (Uniprot 2021). The database search results were loaded onto Scaffold Q+ (Proteome Sciences) for statistical treatment and data visualization. Protein cutoffs were set to a protein threshold of 99.0%, a minimum peptide number of 2, and a peptide threshold of 95%. Differentially expressed proteins (DEPs) between conditions were analyzed using t-tests on normalized spectral values. Proteins with p-values ≤ 0.05 were considered differentially expressed.

### Pathway overrepresentation analysis

Pathway analysis was conducted using Gene Ontology (GO) term overrepresentation analysis for biological processes, implemented with the clusterProfiler package in R. Pathways with a Benjamini Hochberg corrected p-value < 0.05 and a minimum of three proteins were considered significant. The top 15 pathways in response to cannabis smoke versus air were visualized through heatmaps and network plots. Additionally, proteins associated with key pathways were further examined using heatmaps based on their z-scores, generated with the pheatmap R package.

### Blood and lung flow cytometry

Blood was obtained through cardiac puncture and directly stained with conjugated antibodies (Supp. Table 1, Supp. Fig. 1), after which red blood cells were lysed. For lung tissue, an enzyme cocktail consisting of 10 units of DNase, 0.4 units of Collagenase D, and HBSS to a final volume of 2.5 ml was prepared for each set of lungs. Lungs were injected with 1 ml of the enzyme cocktail, tied off, and added to a tube containing an additional 1.5 ml of the enzyme cocktail. Lungs were digested using the gentleMACS™ Octo Dissociator with Heaters (Miltenyi Biotec; Gaithersburg, MD) and strained through a 90-µm mesh. Red blood cells were lysed using RBC Lysis Buffer (BioLegend; San Diego, CA) according to the manufacturer’s instructions and cells were resuspended in PBS. For flow cytometry, single cell suspensions were stained with eBioscience Fixable Viability Dye (Invitrogen; Burlington, ON), followed by incubation with Fc block (BioLegend; San Diego, CA), and finally stained with fluorescent-conjugated primary antibodies (Supp. Table 2, Supp. Fig. 2). Data were acquired on a BD LSR Fortessa™ Cell Analyzer and analyzed using FlowJo software version 10.0 for Macintosh.

### Enzyme-linked immunosorbent assay (ELISA)

Concentration of the THC metabolite 11-Nor-9-carboxy-Δ^9^-tetrahydrocannabinol (THC-COOH) in plasma was analyzed using a direct competitive THC Forensic ELISA kit (NEOGEN®, KY), according to the manufacturer’s instructions, using positive and negative controls for blood as previously described (32). The absorbance was read at 450 nm using an Infinite TECAN (TECAN; Baldwin Park, CA).

### RT-qPCR

Total RNA was extracted using the Aurum Total RNA Mini Kit (Bio-Rad; Saint Laurent, QC) according to the manufacturer’s instructions. RNA quantification was performed on an Infinite TECAN (TECAN; Baldwin Park, CA). Reverse transcription of RNA was carried out using the iScript™ Reverse Transcription Supermix (Bio-Rad, CA). The resulting cDNA template was used to measure mRNA levels. SsoFast™ EvaGreen® (Bio-Rad; Saint Laurent, QC) was used for the PCR. All results were normalized using 18S RNA as the housekeeping gene. Fold change was calculated using the −ΔΔCt method, and results are presented as fold-change normalized to the housekeeping gene. Genes and primer sequences used are listed in Supp. Table 3.

### Multiplex analysis

BALF and plasma were analyzed using multiplex analysis of inflammatory markers (Eve Technologies; Calgary, AB). Cytokines were assessed using the Mouse Cytokine/Chemokine 31-Plex Discovery Assay®.

### Statistical analysis

Statistical analysis was performed using GraphPad Prism v9.0 (GraphPad Software, San Diego, CA). All results are reported as mean ± SEM. ANOVA followed by Sidak’s multiple comparisons test was performed to determine statistical significance unless otherwise indicated.

### Study approval

This study was approved by the McGill University Animal Care Committee and carried out in accordance with the Canadian Council on Animal Care.

## Results

### Cannabis Smoke Exposure Activates AhR in Lung Tissue

To investigate whether the effects of cannabis smoke on the pulmonary system are regulated by the AhR, we exposed *Ahr^+/-^* and *Ahr^-/-^* mice to two cannabis smoke exposures per day for three days. Selecting an appropriate dose is important for the translation of preclinical findings to human health; therefore, we first measured serum levels of the THC metabolite THC-COOH in cannabis-exposed *Ahr^+/-^* and *Ahr^-/-^* mice (Fig. 1A). The exposure regimes used in this study resulted in THC-COOH concentrations of 59.7 ng/ml in *Ahr^+/-^* mice and 61.03 ng/ml in *Ahr^-/-^* mice, levels slightly below those observed in daily cannabis users (>100 ng/ml) (33), making this a moderate exposure regimen. Additionally, the similarity of THC-COOH levels between the *Ahr^+/-^* and *Ahr^-/-^* mice indicates that AhR does not impact THC metabolism in this model. We next assessed whether cannabis smoke activates the AhR in lung tissue by measuring mRNA levels of the well-established AhR target genes *Cyp1a1* and *Cyp1b1* (Fig. 1B-C). *Cyp1a1* and *Cyp1b1* mRNA was significantly elevated in cannabis-exposed *Ahr^+/-^* but not *Ahr^-/-^* mice, demonstrating that cannabis smoke activates the AhR at the exposure level delivered to the mice via inhalation.

**Figure 1.**
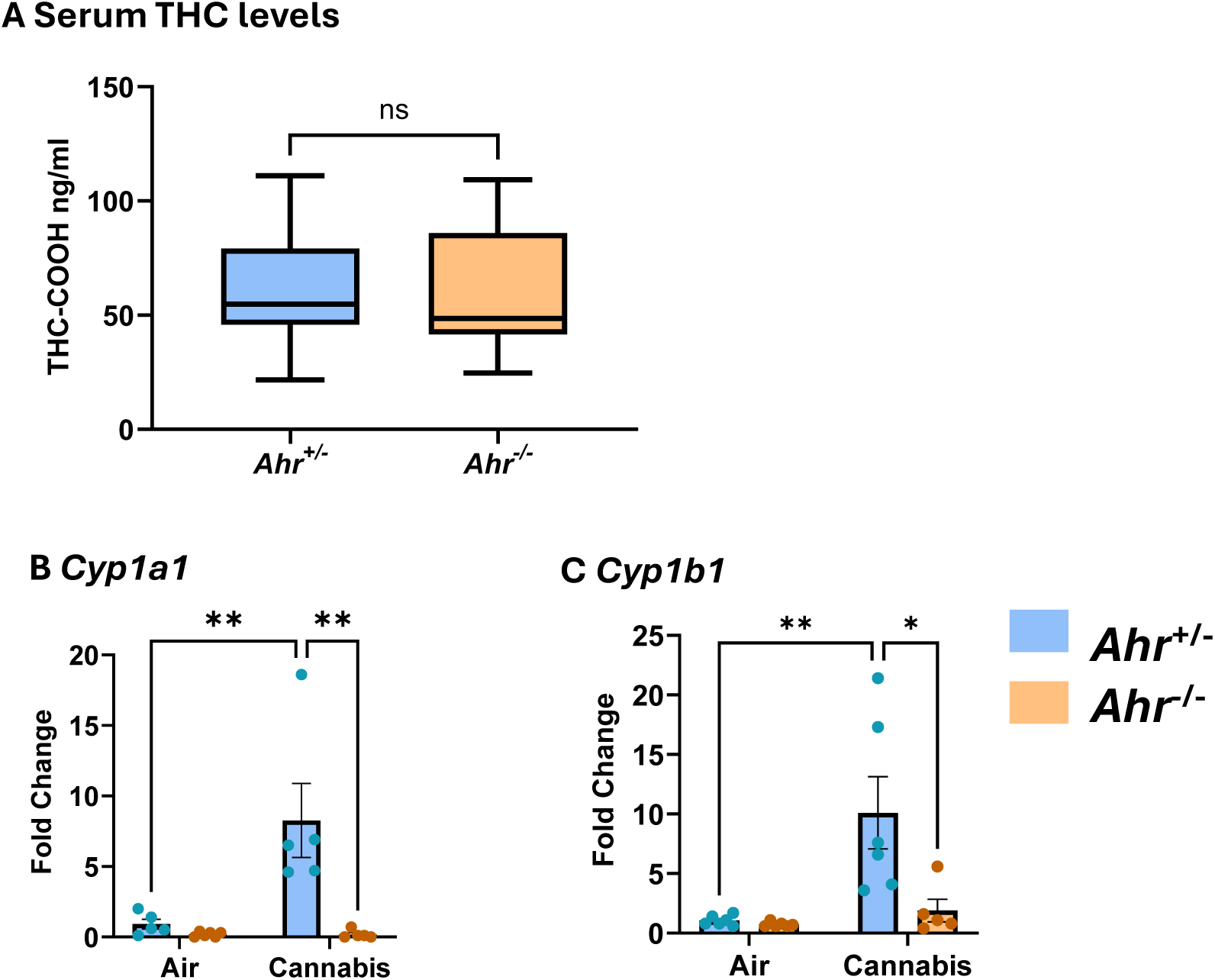
Moderate cannabis smoke exposure activates pulmonary AhR. (A) Serum levels of THC-COOH after acute cannabis smoke exposure in *Ahr^+/-^* and *Ahr^-/-^* mice shown as box plots indicating the mean, minimum, and maximum values. Relative levels of *Cyp1a1* (B) and *Cyp1b1* (C) mRNA in lung tissue after air or cannabis smoke exposure in *Ahr^+/-^* and *Ahr^-/-^* mice. Data represent mean ± SEM, with individual points indicating biological replicates. Statistical significance is shown as *p < 0.05, **p < 0.01.

### Cannabis Smoke Causes Epithelial Sloughing and AhR-Regulated Neutrophilia

Cannabis smoke contains various combustion products, including carcinogens, mutagens, and respiratory toxicants (6). Exposure to these compounds can trigger acute inflammation, primarily mediated by the innate immune system (34). AhR protects against environmental pollutants, but its role in the innate immune response to cannabis smoke is unknown. To address this, we performed a complete profiling of innate immune cell populations in the lung, BALF, and blood of *Ahr^+/-^* and *Ahr^-/-^* mice following cannabis smoke exposure (Fig. 2A).

**Figure 2.**
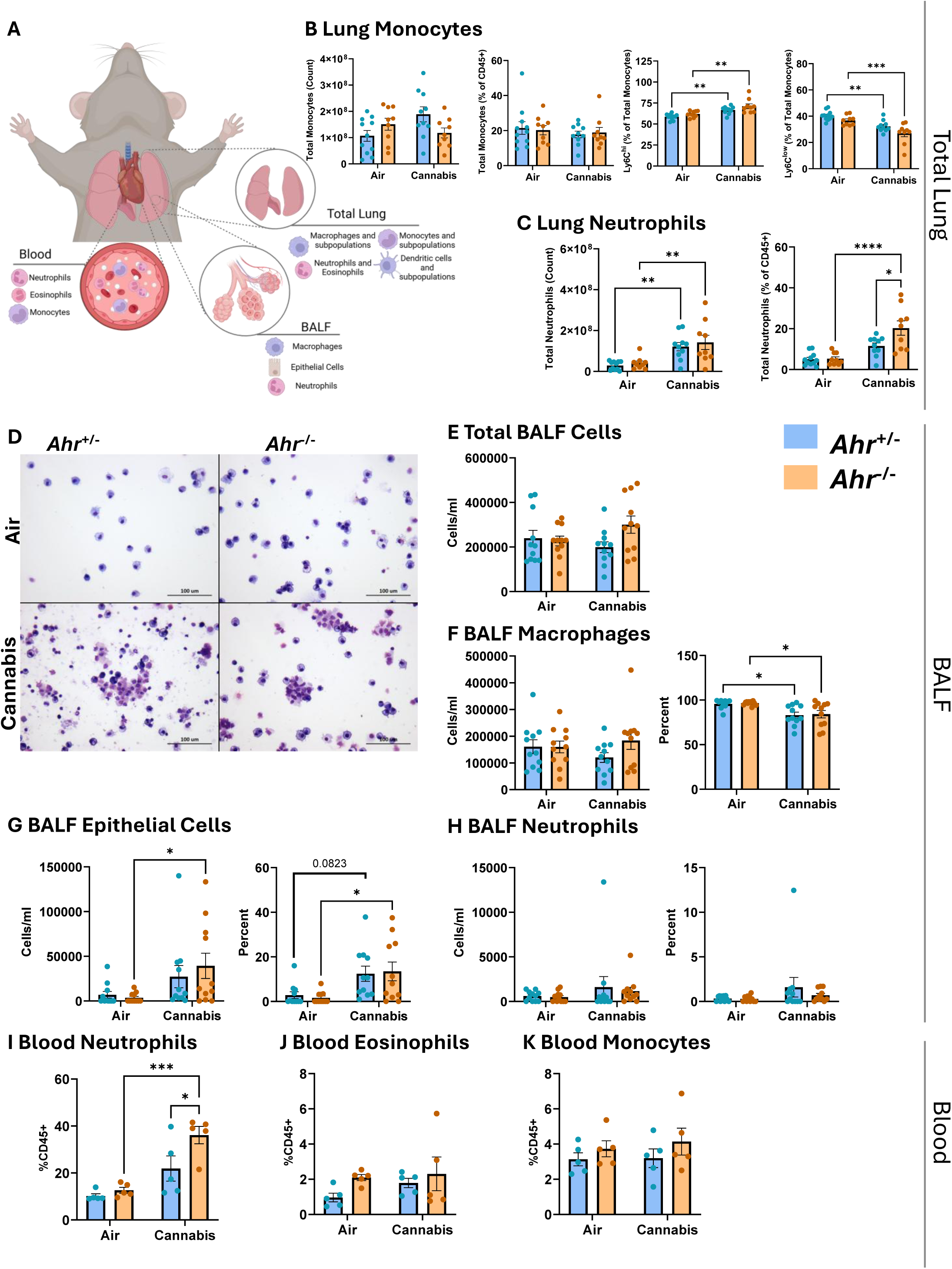
Cannabis smoke alters innate immune cell populations in the lung, airspaces, and blood. (A) Schematic representation of immune cell populations assessed in the lung, airspaces, and blood. (B-C) Pulmonary monocytes and neutrophils, including total counts and percentages of CD45^+^ cells. (D) Representative images of BALF cells from *Ahr^+/-^* and *Ahr^-/-^* mice after air or cannabis smoke exposure. Scale bars represent 100 µm. (E-H) Differential cell counts in BALF, including total BALF cells (E), macrophages (F), epithelial cells (G), and neutrophils (H), with total counts and percentages shown. (I-K) Innate immune cell populations in blood, including neutrophils (I), eosinophils (J), and monocytes (K). Data represent mean ± SEM, with individual points indicating biological replicates. Statistical significance is denoted as *p < 0.05, **p < 0.01, ***p < 0.001, and ****p < 0.0001.

In the total lung, cannabis smoke did not affect macrophage, eosinophil, or dendritic cell populations (Supp. Fig. 3). Monocytes, which are recruited to the lungs during inflammation and can differentiate into macrophages, remained unchanged (Fig. 2B); however, cannabis smoke exposure caused a shift in monocyte subpopulations. Here, there was an increase in Ly6C^hi^ monocytes and a decrease in Ly6C^low^ monocytes in both *Ahr^+/-^* and *Ahr^-/-^* mice, suggesting an emerging inflammatory response. Consistent with this possibility, cannabis smoke significantly increased pulmonary neutrophil counts, with a higher percentage of neutrophils in *Ahr^-/-^* mice compared to *Ahr^+/-^* mice (Fig. 2C). This finding aligns with previous studies showing that AhR inhibits neutrophil recruitment to the lungs in response to cigarette smoke (28).

We next investigated whether cannabis smoke exposure recruits immune cells to the airways by assessing the cellular composition of BALF from *Ahr^+/-^* and *Ahr^-/-^* mice after acute exposure to cannabis smoke or air (Fig. 2D). The total number of cells in the BALF was similar across all experimental groups (Fig. 2E); however, we observed differences in the cellular makeup of the BALF compartment. Macrophages were the predominant cell type, and while their total numbers remained unchanged, cannabis smoke exposure reduced the percentage of macrophages in the BALF in both genotypes (Fig. 2F). This reduction in frequency of macrophages was accompanied by a rise in epithelial cells present in BALF after cannabis smoke exposure, indicating epithelial sloughing (Fig. 2G). Although both genotypes showed an increase in epithelial cells, the change was significant only in *Ahr^-/-^* mice, while smoke-exposed *Ahr^+/-^* did not reach significance (p=0.082). We next investigated whether the increase in neutrophils within the lung tissue was also reflected in the airways. However, cannabis smoke did not elevate neutrophils in BALF (Fig. 2H), suggesting that neutrophils recruited in response to cannabis smoke remain in the lung interstitium without entering the airways. Thus, cannabis smoke leads to sloughing of the respiratory epithelium, with *Ahr^-/-^* mice showing heightened susceptibility to barrier disruption.

Finally, we analyzed the effects of cannabis smoke on systemic inflammation by examining immune cells present in the blood. Neutrophils were significantly increased in the blood of *Ahr^-/-^* mice in response to cannabis smoke (Fig. 2I), whereas the percent of eosinophils and monocytes remained unchanged (Fig. 2J-K). These findings suggest that AhR not only plays a role in protecting the lungs from inflammation but can also help safeguard other organs by suppressing systemic inflammation.

### Cannabis Smoke Triggers Local and Systemic Inflammatory Cytokines

We also evaluated cytokine levels in both the BALF and blood to assess local and systemic inflammatory signals following cannabis smoke exposure. Beyond immune cell recruitment, cytokines regulate immune activation, tissue remodeling, and communicate inflammatory signals to organs beyond the lungs. In the BALF, cannabis smoke exposure significantly increased VEGF and LIF levels in both *Ahr^+/-^* and *Ahr^-/-^* mice (Fig. 3A), which suggests activation of tissue remodeling and repair pathways consistent with the epithelial damage observed in Figure 2. Additionally, there was an AhR-independent upregulation of interleukin (IL)-6, a pleiotropic pro-inflammatory cytokine involved in immune cell recruitment and acute-phase protein production, by cannabis smoke in BALF (Fig. 3B). Cannabis smoke also increased the eosinophil-related cytokines eotaxin and IL-5 (Fig. 3C), indicating a shift toward an eosinophil-mediated immune response. Although the increase in eotaxin was independent of AhR, only *Ahr^+/-^* mice had an increase in IL-5 in response to cannabis smoke, suggesting that AhR is needed for its induction in the BALF. Interestingly, cannabis smoke reduced the T-cell-associated cytokines IL-2 and RANTES only in cannabis smoke-exposed *Ahr^-/-^* mice BALF (Fig. 3D), suggesting AhR is needed to maintain the levels of these cytokines. BALF cytokines unaffected by cannabis smoke are listed in Supplemental Table 4.

**Figure 3.**
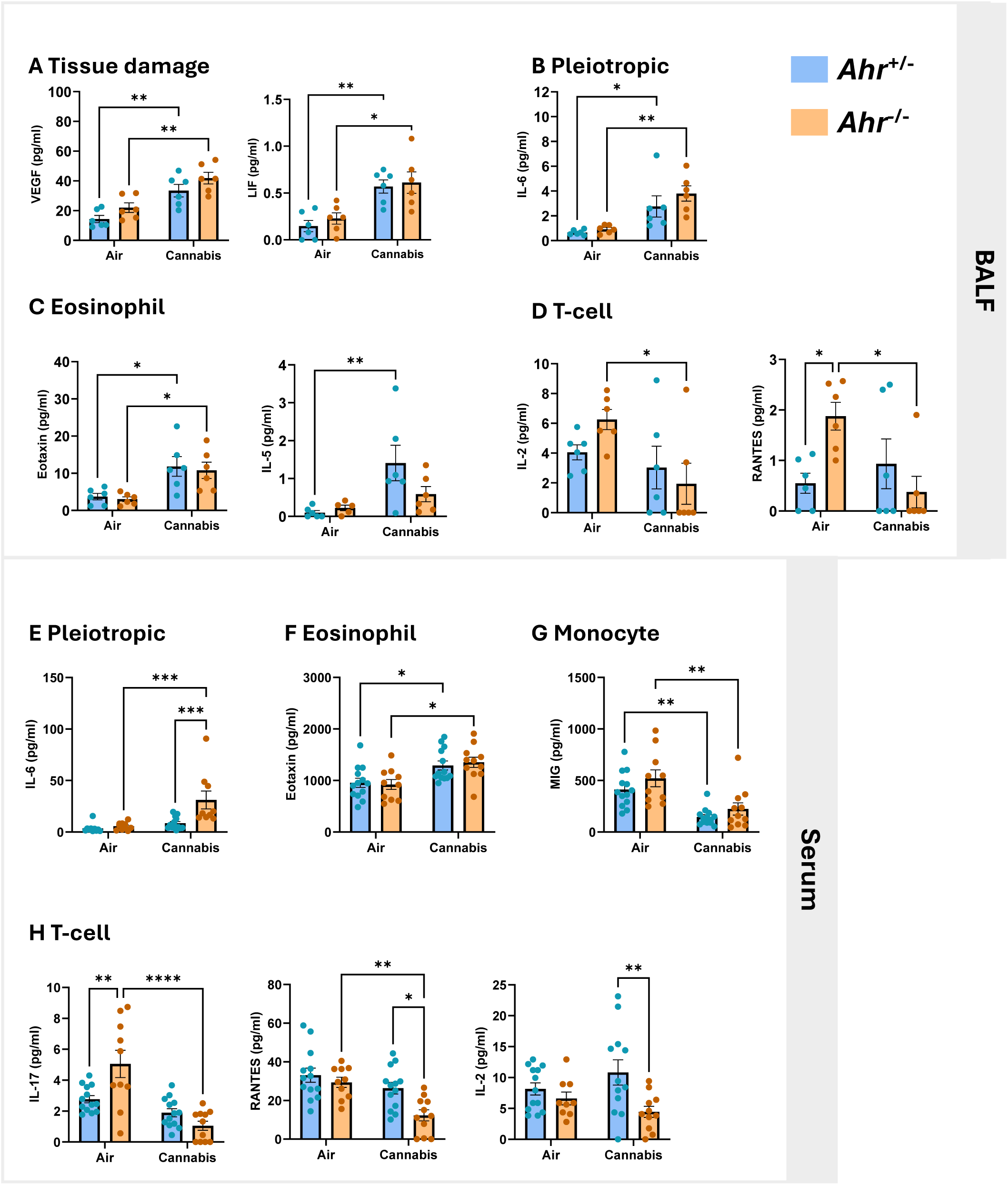
Cannabis smoke induces cytokine expression in BALF and serum. Cytokine concentrations were measured in BALF (A-D) and serum (E-H) from *Ahr^+/-^* and *Ahr^-/-^* mice following exposure to air or cannabis smoke. (A) Tissue damage-related cytokines VEGF and LIF. (B) Pleiotropic cytokine IL-6. (C) Eosinophil-related cytokines eotaxin and IL-5. (D) T-cell-related cytokines IL-2 and RANTES. (E) Systemic IL-6 levels in serum. (F) Serum eosinophil-related cytokine eotaxin. (G) Serum monocyte-related cytokine MIG. (H) Serum T-cell-related cytokines IL-17, RANTES, and IL-2. Data are presented as mean ± SEM, with individual points representing biological replicates. Statistical significance is indicated as *p < 0.05, **p < 0.01, ***p < 0.001, and ****p < 0.0001.

Beyond the local lung environment, cannabis smoke also altered systemic inflammation. There was a significant increase in IL-6 only in *Ahr^-/-^* mice exposed to cannabis smoke, which may explain the elevated neutrophils in the blood of *Ahr^-/-^* mice (Fig. 3E). Eotaxin levels were elevated by cannabis smoke independent of AhR (Fig. 3F), suggesting eosinophil activation. Conversely, levels of the monocyte-associated cytokine MIG were reduced by cannabis smoke exposure in the blood of both *Ahr^+/-^* and *Ahr^-/-^* mice (Fig. 3G). Consistent with findings in the BALF, the T-cell-associated cytokines IL-17, RANTES, and IL-2 were significantly reduced in the blood of *Ahr^-/-^* mice but remained unchanged in *Ahr^+/-^* mice (Fig. 3H), further supporting the role of AhR in maintaining adaptive immune function following cannabis smoke exposure. Serum cytokines unaffected by cannabis smoke are listed in Supplemental Table 5. Overall, these results demonstrate that cannabis smoke triggers both local and systemic inflammatory responses, characterized by increased cytokines involved in tissue repair and innate immune cell recruitment.

### Characterization of the Pulmonary Proteome in Response to Cannabis Smoke

The respiratory system produces a complex secretome that contributes to lung homeostasis and immune responses. This secretome includes EVs and soluble proteins which are secreted into the airways and can serve as messengers of intercellular communication. To gain a deeper understanding of how cannabis smoke and AhR intersect to influence the respiratory proteome, we performed proteomic analysis to comprehensively profile the lung tissue, EVs, and BALF (Fig. 4A). This comprehensive approach allowed for assessment of both the cellular and extracellular proteomes, providing insight into how cannabis smoke impacts lung secretions and intercellular signaling. First, using TEM, we observed heterogeneous populations of EVs across all experimental conditions (Fig. 4B). The EV population exhibited broad size variation, with most vesicles measuring <200 nm, suggesting the EVs primarily consist of microvesicles and exosomes, rather than apoptotic bodies (Fig. 4C). Neither cannabis smoke exposure nor AhR status significantly altered the overall concentration of EVs released into the airways (Fig. 4D). These findings suggest that the total abundance of EVs remains stable following cannabis smoke exposure.

**Figure 4.**
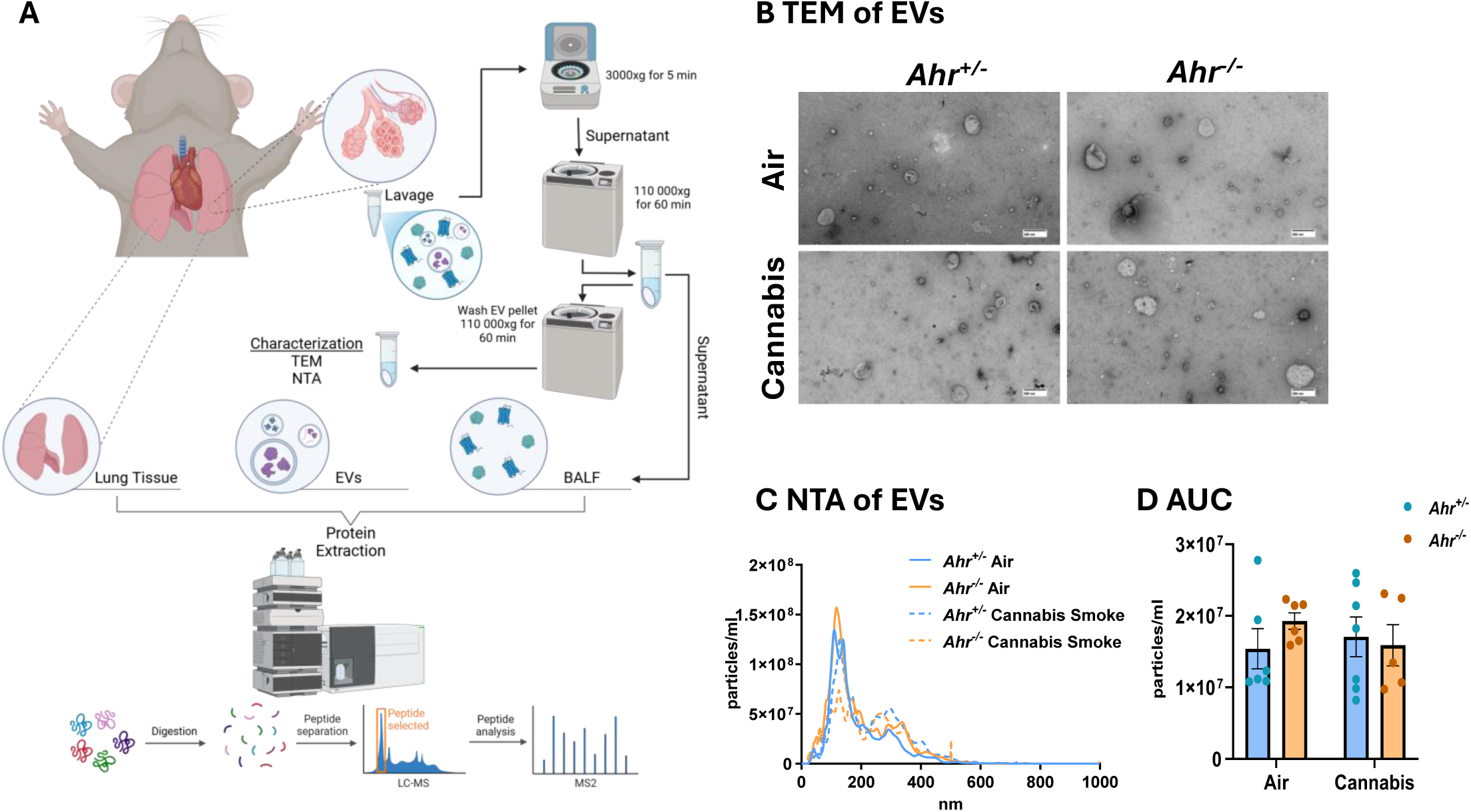
Characterization of pulmonary compartments for proteomic analysis. (A) Schematic representation of the experimental workflow used to define and isolate lung tissue, EVs, and BALF for proteomic analysis. (B-D) Characterization of EVs including (B) TEM images of EVs from *Ahr^+/-^* and *Ahr^-/-^* mice following exposure to air or cannabis smoke. Scale bar represents 200 nm. (C) NTA of EVs, showing particle size distribution across 0–1000 nm. (D) Area under the curve (AUC) analysis of total EV concentration in BALF. Data represent mean ± SEM, with individual points indicating biological replicates.

To investigate how cannabis smoke and AhR influence proteomic profiles, bottom-up LC-MS/MS analysis was performed on each compartment. DEPs and pathways were analyzed to identify features across compartments, illustrating the unique and collective role of these compartments in the pulmonary response to cannabis smoke. First, we profiled pathways that were upregulated by cannabis smoke in *Ahr^+/-^* mice, thereby reflecting how cannabis smoke augments the proteomic response. Here, proteins within lung tissue and EVs exhibited the greatest number of upregulated proteins, with 71 proteins unique to the lung and 57 unique to EVs (Fig. 5A, Supp. Table 6). Among the three pulmonary compartments, EVs and BALF shared the largest overlap, with 13 upregulated DEPs common between them. GO term enrichment was performed on the upregulated proteins in each compartment and this analysis revealed distinct as well as shared pathway activation patterns (Fig. 5B, Supp. Table 7). In *Ahr^+/-^* mice, the lung tissue displayed the most proteomic changes, with 157 exclusively upregulated pathways. To understand the interplay between compartments, we performed network analysis on the top 15 upregulated pathways from lung, EVs, and BALF (Fig. 5C). EV and BALF pathways clustered together, and were predominantly associated with hemostasis and coagulation, suggesting a similar response to cannabis smoke. In contrast, lung-specific pathways formed distinct clusters, largely related to metabolism and detoxification. Negative regulation of proteolysis and peptidase activity was also specific to the BALF compartment. Collectively, these data demonstrate that each compartment plays a distinct role in the pulmonary response to cannabis smoke exposure, with the networks between EV and BALF pathways suggesting a complementary function in extracellular signaling.

**Figure 5.**
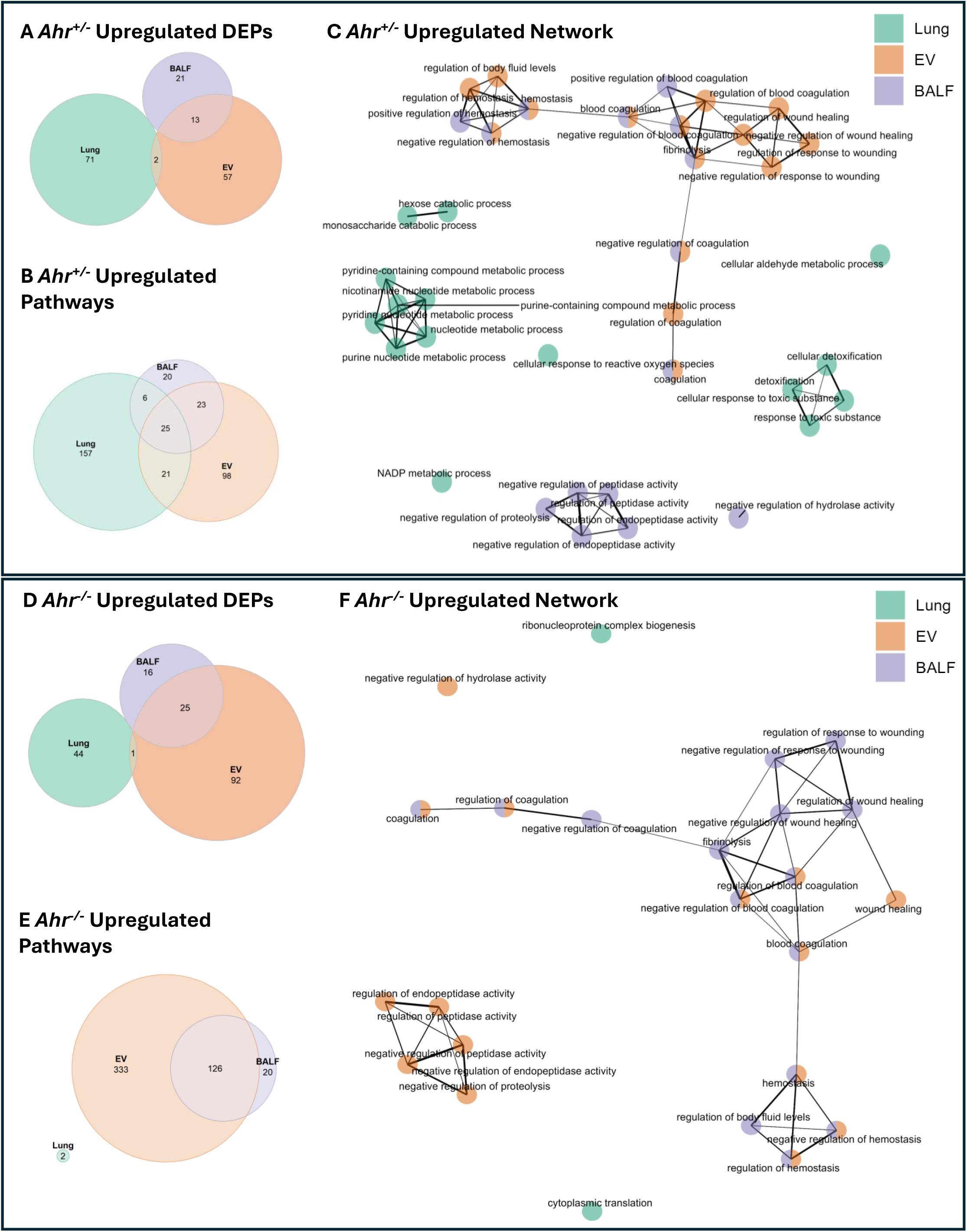
Overlapping and unique upregulated DEPs and pathways across lung compartments in response to cannabis smoke. Venn diagram of upregulated DEPs (A) and pathways (B) in *Ahr^+/-^* mice exposed to cannabis smoke versus air. (C) Network analysis of upregulated pathways in *Ahr^+/-^*, with each node representing biological processes enriched in lung (green), EVs (orange), or BALF (purple). Venn diagram of upregulated DEPs (D) and pathways (E) in *Ahr^-/-^* mice exposed to cannabis smoke versus air. (F) Network analysis of upregulated pathways in *Ahr^-/-^*, with each node representing biological processes enriched in lung (green), EVs (orange), or BALF (purple).

Next, we evaluated proteins in cannabis smoke-exposed *Ahr^-/-^* mice. Here, cannabis smoke had the greatest impact on the EV compartment, with 92 unique upregulated proteins (Fig. 5D, Supp. Table 8). The notable overlap of upregulated proteins between EVs and BALF, particularly the presence of annexin A4 (Anxa4), annexin A5 (Anxa5), and annexin A11 (Anxa11), suggests a coordinated extracellular response to cannabis smoke exposure caused by AhR deficiency. The increased presence of these proteins indicates that cannabis smoke could affect cell growth, survival, and membrane repair in lung tissue (35). GO term enrichment analysis revealed that the EVs and BALF compartments contained the most enriched pathways, whereas the lung compartment of these *Ahr^-/-^* mice had only two upregulated pathways following cannabis smoke exposure (Fig. 5E, Supp. Table 9). Network analysis of the top 15 upregulated pathways in each compartment of *Ahr^-/-^* mice showed that EV and BALF pathways clustering together and were related to hemostasis and coagulation (Fig. 5F). While negative regulation of proteolysis and peptidase activity was exclusive to BALF in *Ahr^+/-^* mice, this process shifted to the EV compartment in *Ahr^-/-^* mice, suggesting that AhR may influence compartment-specific responses to proteolytic regulation. Collectively, these data demonstrate that each compartment contributes uniquely to the pulmonary response to cannabis smoke exposure. The presence of AhR appears to facilitate a robust proteomic response in lung tissue, while simultaneously restraining the upregulated response in EVs, suggesting a regulatory role in extracellular signaling. In *Ahr^-/-^* mice, the lack of AhR-driven tissue response may be compensated by changes in the secretome, particularly in EVs and BALF, highlighting a potential mechanism by which the lung adapts to environmental stressors in the absence of AhR.

Our proteomic analysis also revealed downregulation in numerous proteins and pathways caused by cannabis smoke that were both AhR-dependent and AhR-independent. Downregulated proteins in *Ahr^+/-^* mice were also predominantly found in the lung tissue, with 329 unique DEPs (Fig. 6A, Supp. Table 10). GO term enrichment revealed that lung tissue exhibited the greatest number of downregulated pathways (Fig. 6B). Furthermore, lung and BALF had the most shared downregulated pathways which were primarily related to remodeling processes (Supp. Table. 11). Network analysis of the top 15 pathways from lung, EVs, and BALF revealed small pathway clusters that were primarily compartment specific, with little overlap between them (Fig. 6C). The largest cluster, found in lung tissue and BALF, was associated with actin and cytoskeletal organization. BALF-specific pathways were related to epithelial cell migration. Finally, EV and BALF shared a cluster of pathways related to heat and temperature stimulus response, suggesting a coordinated reaction to external stress.

**Figure 6.**
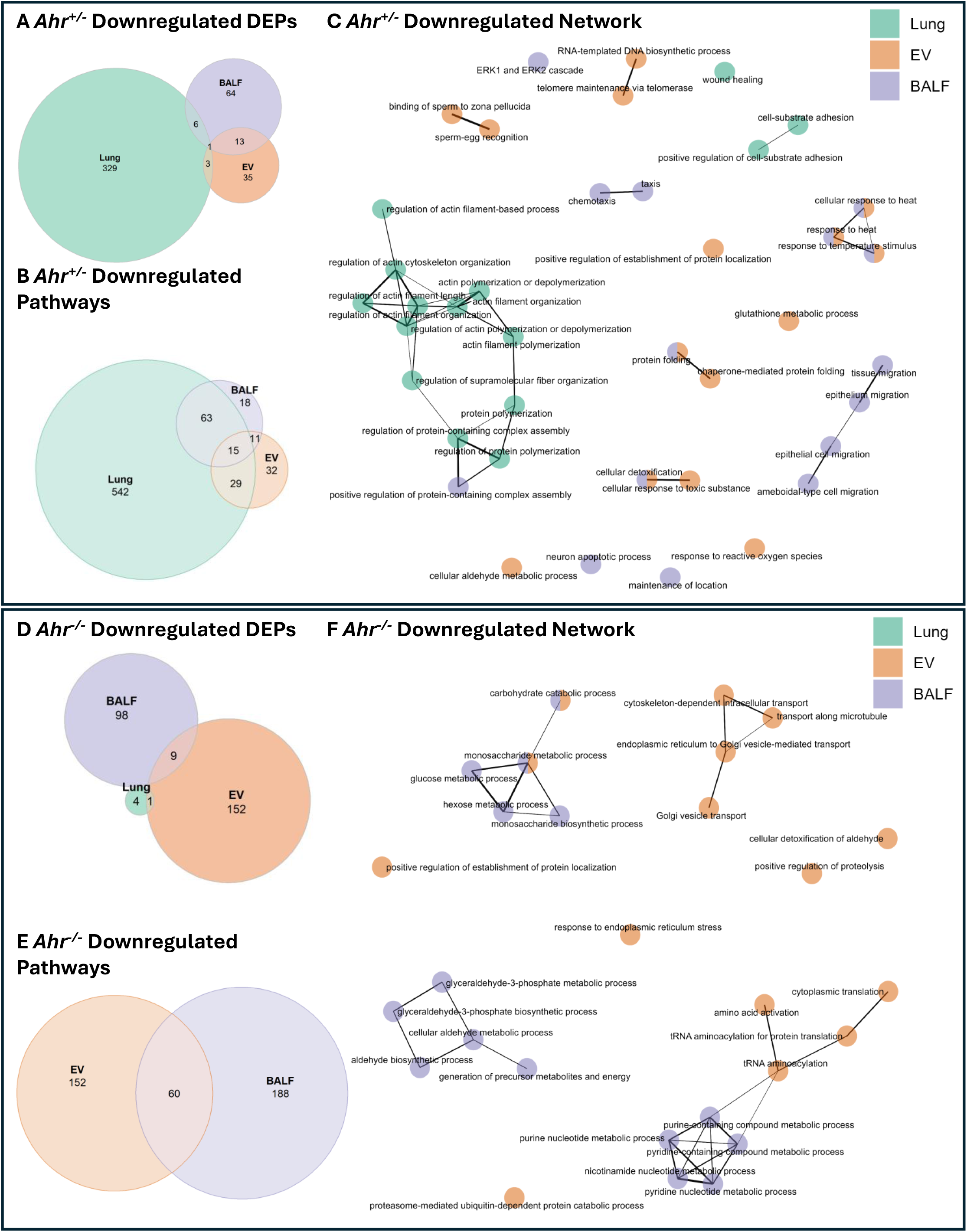
Overlapping and unique downregulated DEPs and pathways across lung compartments in response to cannabis smoke. Venn diagram of downregulated DEPs (A) and pathways (B) in *Ahr^+/-^* mice exposed to cannabis smoke versus air. (C) Network analysis of upregulated pathways in *Ahr^+/-^* mice, with each node representing biological processes enriched in lung (green), EVs (orange), or BALF (purple). Venn diagram of downregulated DEPs (D) and pathways (E) in *Ahr^-/-^*mice exposed to cannabis smoke versus air. (F) Network analysis of upregulated pathways in *Ahr^-/-^*, with each node representing biological processes enriched in lung (green), EVs (orange), or BALF (purple).

In contrast, there was little downregulation of proteins in lung tissue from cannabis smoke-exposed *Ahr^-/-^* mice. Instead, nearly all downregulation of proteins occurred in EVs and BALF (Fig. 6D, Supp. Table 12), highlighting a potential role of the secretome in compensating for the diminished lung tissue response. Pathway enrichment analysis showed that all downregulated pathways were restricted to EVs and BALF, with 60 pathways shared between them (Fig. 6E, Supp. Table 13). Network analysis further emphasized the coordinated response between EVs and BALF, particularly in metabolic process regulation (Fig. 6F), highlighting their interconnected role in the pulmonary secretome. Collectively, these findings demonstrate the distinct yet correlated response of each compartment to cannabis smoke exposure, emphasizing the importance of studying the secretome alongside the cellular response of the pulmonary system.

### AhR Activation Induces Lung Detoxification Pathways While Downregulating Structural Integrity in Response to Cannabis Smoke

Our analysis above showed that cannabis smoke elicits significant and distinct processes within the respiratory system, with some of the largest alterations seeming to occur in an AhR-dependent manner. Therefore, we next compared the proteomic response of cannabis smoke versus air for *Ahr^+/-^* and *Ahr^-/-^* mice for each pulmonary compartment, beginning with the lung tissue. In the lungs of *Ahr^+/-^* mice, cannabis smoke exposure resulted in a greater number of DEPs than in lung from *Ahr^-/-^* mice (Supp. Fig. 4A). While cannabis smoke primarily suppressed DEPs in the lungs of *Ahr^+/-^* mice, it had the opposite effect in *Ahr^-/-^* mice, where most DEPs were upregulated (Supp. Fig. 4B-D). Focusing on the upregulated proteins, lung tissue from *Ahr^+/-^* and *Ahr^-/-^* mice shared only three overlapping proteins following cannabis smoke exposure (Fig. 7A, Supp. Table 14). When considering both genotypes together, all of the top 15 upregulated pathways within lung tissue in response to cannabis smoke were exclusive to *Ahr^+/-^* mice (Fig. 7B, Supp. Table 15). To assess how these pathways were related, we performed clustering analysis, which revealed two major clusters: detoxification and nucleotide metabolism, in which all the proteins were only differentially expressed in the *Ahr^+/-^* mice (Fig. 7C, Supp. Fig. 5A). To further examine the protein-level changes underlying these pathways, we identified the top 15 proteins from each cluster (Fig. 7D-E). In the detoxification cluster, proteins such as Aldh1a7, Cat, Prdx1, Aldh1a1, and Cyp2f2 were expressed at lower levels in lung tissue from *Ahr^+/-^* mice at baseline but were induced in response to cannabis smoke only in *Ahr^+/-^* mice (Fig. 7D). These proteins play key roles in oxidative stress defense, aldehyde detoxification, and phase I/II metabolism, suggesting that AhR activation enhances the ability of the lungs to metabolize reactive byproducts of cannabis smoke. In the nucleotide metabolism cluster, proteins including Eno1, Aldh1a7, Gapdh, Vcp, Fmo1, and Idh2 exhibited a similar pattern of low baseline expression in *Ahr^+/-^* lungs and induction following cannabis smoke exposure (Fig. 7E). These proteins regulate purine and pyrimidine metabolism, nucleotide biosynthesis, and redox balance, indicating that AhR may coordinate metabolic shifts to support cellular adaptation to cannabis smoke. Overall, these findings suggest that AhR signaling is necessary for the metabolic adaptation of lung tissue to cannabis smoke, specifically through the activation of detoxification and nucleotide metabolism pathways.

**Figure 7.**
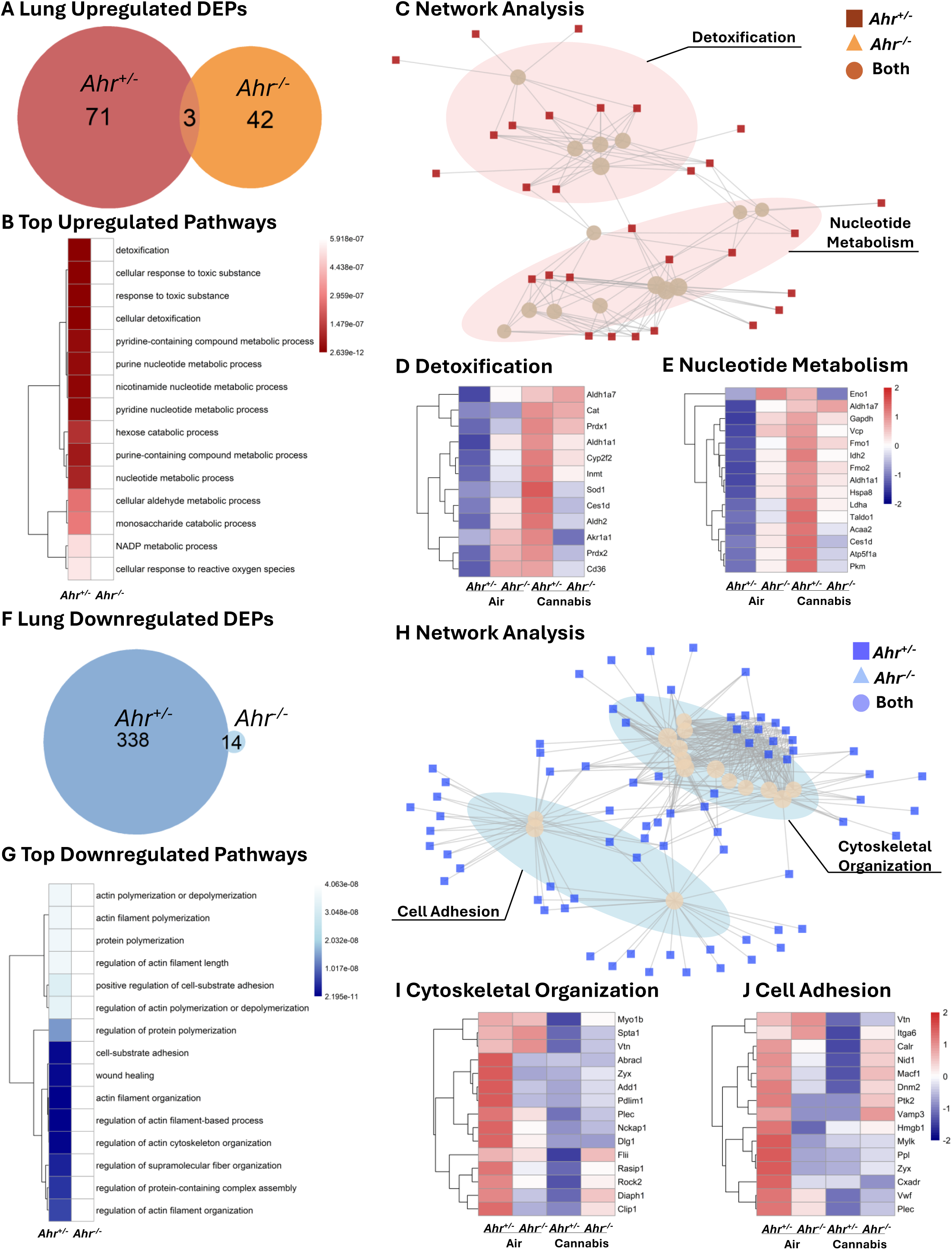
Pathway analysis of DEPs in lung tissue of *Ahr^+/-^* and *Ahr^-/-^* mice exposed to cannabis smoke versus air. (A) Venn diagram showing the overlap of upregulated DEPs for *Ahr^+/-^* and *Ahr^-/-^* lung tissue after cannabis smoke exposure versus air. (B) Top 15 upregulated pathways induced by cannabis smoke exposure versus air in *Ahr^+/-^* and *Ahr^-/-^* mice. (C) Network clustering analysis of upregulated pathways. (D-E) Heatmaps of top 15 proteins in each cluster including detoxification (D) and nucleotide metabolism (E). (F) Venn diagram of downregulated DEPs for *Ahr^+/-^* and *Ahr^-/-^* lung tissue after cannabis smoke exposure versus air. (G) Top 15 downregulated pathways in response to cannabis smoke exposure in *Ahr^+/-^* and *Ahr^-/-^* mice. (H) Network analysis of downregulated pathways. (I-J) Heatmaps of top 15 proteins in the cytoskeletal organization (I) and cell adhesion (J) clusters. Heatmaps represent z-scores of protein expression across conditions.

Protein downregulation in lung tissue after cannabis exposure was also most prevalent in *Ahr^+/-^* mice (Fig. 7F, Supp. Table 16) with a decrease in the expression of 339 DEPs compared to 5 DEPs in *Ahr^-/-^* mice. The top 15 downregulated pathways in response to cannabis smoke were primarily related to actin filament organization, cytoskeletal regulation, and cell adhesion, and these pathways were significantly suppressed only in *Ahr^+/-^* mice (Fig. 7G, Supp. Table 17). Network analysis further showed that these pathways clustered into two major groups: cytoskeletal organization and cell adhesion (Fig. 7H, Supp. Fig. 5B). To examine the underlying protein-level changes, we identified the top 15 proteins from each pathway cluster (Fig. 7I-J). In the cytoskeletal organization cluster, proteins such as Myo1b, Spta1, Vtn, Abracl, Zyx, and Add1 were highly expressed in *Ahr^+/-^* lungs at baseline but showed reduced expression following cannabis exposure while *Ahr^-/-^* mice exhibited consistently low expression of these proteins, unaffected by cannabis smoke (Fig. 7I). These proteins regulate actin polymerization, filament stability, and cytoskeletal remodeling, suggesting that AhR regulates cytoskeletal integrity in response to environmental exposures. Similarly, in the cell adhesion cluster, proteins such as Cxadr, Itga6, Calr, Nid1, and Macf1 followed the same pattern of high baseline expression in *Ahr^+/-^* lungs and suppression after cannabis exposure (Fig. 7J, Supp. Fig. 6A-C). These proteins are involved in cell-cell and cell-matrix adhesion, extracellular matrix (ECM) interactions, and mechanical stability of epithelial cells. Their suppression suggests that AhR-dependent signaling regulates lung tissue integrity and adhesion properties following cannabis exposure. Overall, these findings indicate that AhR is essential for maintaining cytoskeletal stability and cell adhesion in the lung at baseline, but its activation in response to cannabis exposure leads to the suppression of these pathways. This suppression may contribute to tissue remodeling and impaired barrier function.

### AhR Controls EV Cargo Involving Proteostasis and Antioxidant Defenses in Response to Cannabis Smoke

We next compared the protein cargo of EVs from *Ahr^+/-^* and *Ahr^-/-^* mice exposed to air or cannabis smoke. In this lung compartment, *Ahr^-/-^* mice exhibited a larger proteomic response to cannabis smoke (Supp. Fig. 7A). While *Ahr^+/-^* mice had more upregulated than downregulated DEPs, *Ahr^-/-^* mice showed the opposite pattern (Supp. Fig. 7B-D). When examining the upregulated proteomic cargo in the EVs caused by cannabis smoke, pulmonary EVs from *Ahr^-/-^* mice contained more upregulated DEPs than *Ahr^+/-^* EVs, and the two genotypes shared 32 overlapping DEPs (Fig. 8A, Supp. Table 18). Pathway analysis revealed that EVs from both *Ahr^+/-^* and *Ahr^-/-^* mice exhibited similar upregulated responses to cannabis smoke, with many enriched pathways related to coagulation (Fig. 8B, Supp. Table 19), which is suggestive of vascular leakage, a process often associated with tissue damage or inflammation (36). These pathways included shared proteins such as Klkb1, F2, and the fibrinogen chains Fga, Fgb, and Fgg (Fig. 8C-D, Supp. Fig. 8A). The increased expression of coagulation-related proteins observed in EVs from both *Ahr^+/-^* and *Ahr^-/-^* mice suggests the effects of cannabis smoke on these pathways are AhR-independent.

**Figure 8.**
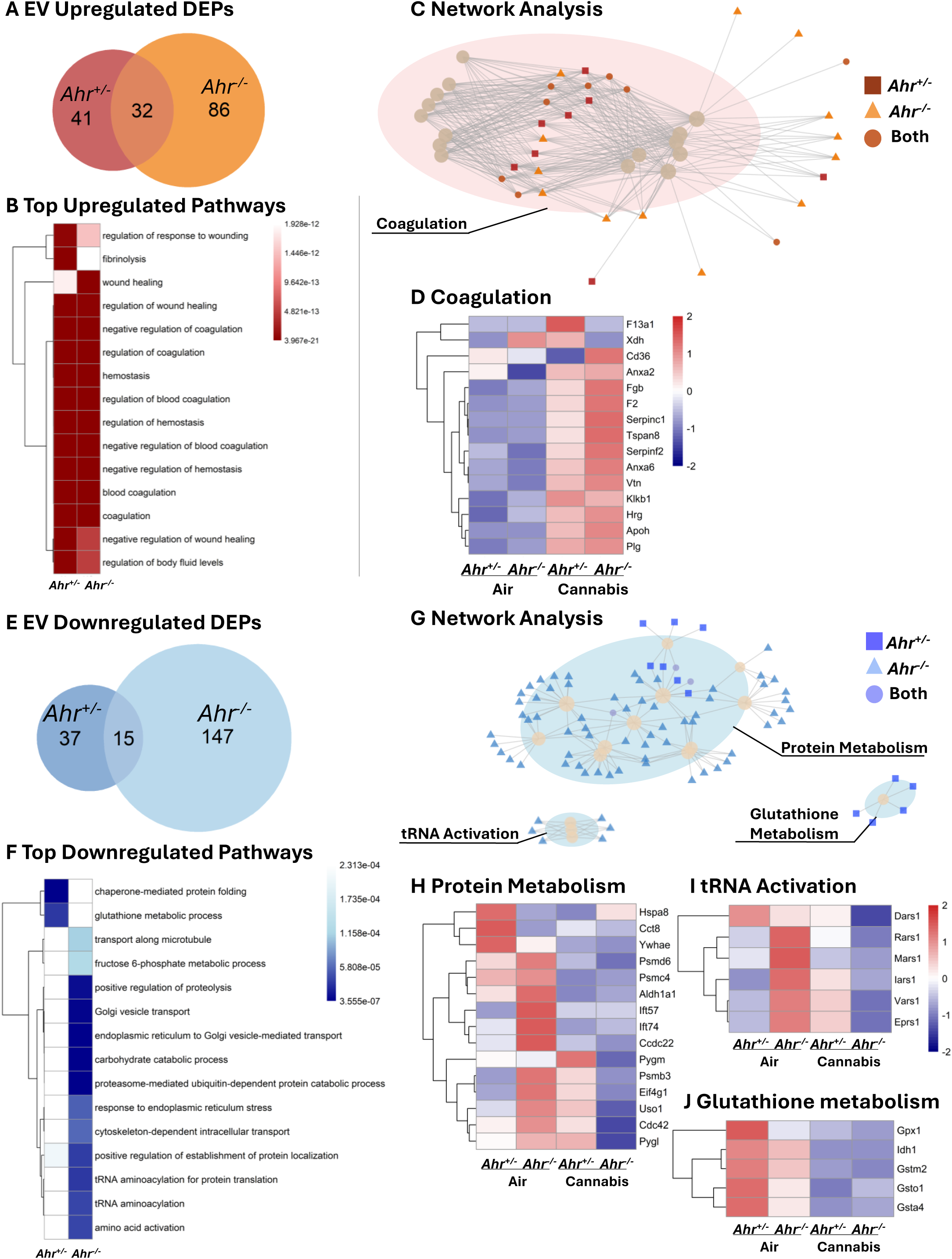
Pathway analysis of DEPs in EVs of *Ahr^+/-^* and *Ahr^-/-^* mice exposed to cannabis smoke versus air. (A) Venn diagram showing the overlap of upregulated DEPs for *Ahr^+/-^* and *Ahr^-/-^* EVs after cannabis smoke exposure versus air. (B) Top 15 upregulated pathways in *Ahr^+/-^* and *Ahr^-/-^* EVs in response to cannabis smoke exposure. (C) Network analysis of upregulated pathways. (D) Heatmap of top 15 coagulation-related proteins. (E) Venn diagram of downregulated DEPs for *Ahr^+/-^* and *Ahr^-/-^* EVs after cannabis smoke exposure versus air. (F) Top 15 downregulated pathways in response to cannabis smoke exposure. (G) Network analysis of downregulated pathways, highlighting clusters related to protein metabolism, tRNA activation, and glutathione metabolism. (H-J) Heatmaps of top 15 proteins in the protein metabolism (H), tRNA activation (I), and glutathione metabolism (J) clusters, showing expression changes in response to cannabis smoke exposure. Heatmaps represent z-scores of protein expression across conditions.

While the upregulated proteomic response revealed similarities between the two genotypes, the downregulated response exhibited more pronounced differences. EVs from *Ahr^-/-^* mice had more downregulated DEPs in response to cannabis smoke, with the two genotypes sharing 15 overlapping DEPs (Fig. 8E, Supp. Table 20). The most significant downregulated EV pathways in response to cannabis smoke were primarily observed in *Ahr^-/-^* mice. (Fig. 8F, Supp. Table 21). Network analysis identified three major pathway clusters: protein metabolism, tRNA activation, and glutathione (GSH) metabolism (Fig. 8G, Supp. Fig. 8B). Protein metabolism-related DEPs generally decreased after cannabis smoke exposure, with a greater number downregulated in EVs from *Ahr^-/-^* mice (Fig. 8H). Notably, proteasome subunits (Psmb7, Psm6, Psmd1, Psmd2, Psmd6, Psmd7, Psmd11, and Psmd13) were significantly reduced only in *Ahr^-/-^* mice (Supp. Fig. 9A-I), suggesting that the AhR maintains proteostasis and protein turnover in EVs in response to cannabis smoke. tRNA activation proteins were highest in EVs from *Ahr^-/-^* mice at baseline, and cannabis smoke reduced their expression exclusively in *Ahr^-/-^* mice, while *Ahr^+/-^* EVs showed a slight increase (Fig. 8I). Lastly, GSH metabolism proteins were highest in EVs from *Ahr^+/-^* mice at baseline but decreased following cannabis exposure in both *Ahr^+/-^* and *Ahr^-/-^* mice (Fig. 8J). This suggests that AhR regulates antioxidant defenses under baseline conditions, but cannabis smoke disrupts redox homeostasis regardless of AhR status. Overall, these findings demonstrate the role of AhR in regulating EV cargo, specifically by maintaining protein turnover and oxidative stress responses following cannabis exposure.

### AhR Modulates Coagulation, Protease Activity, and Metabolic Stability in BALF Following Cannabis Smoke Exposure

The final pulmonary compartment we investigated was the cell-free, EV-free BALF, which contains extracellular proteins released from cells lining the airspaces of lungs. In response to cannabis smoke, BALF from *Ahr^-/-^* mice had more DEPs than *Ahr^+/-^* BALF (Supp. Fig. 10A). In both genotypes, DEPs were predominantly downregulated following cannabis smoke exposure (Supp. Fig. 10B-D), indicating a suppressive effect of cannabis on extracellular protein expression independent of the AhR. Upregulated DEPs were more abundant in the BALF of *Ahr^-/-^* mice, which also shared 20 upregulated DEPs with *Ahr^+/-^* BALF (Fig. 9A, Supp. Table 22). The top upregulated pathways in response to cannabis smoke showed significant overlap between the two genotypes (Fig. 9B, Supp. Table 23). Network analysis grouped these pathways into two main clusters: coagulation and protease regulation (Fig. 9C, Supp. Fig. 11A). In the coagulation cluster, proteins such as Fgb, Fga, Fgg, and Kng1 were elevated in both genotypes following cannabis smoke exposure but were higher in BALF from *Ahr^-/-^* mice (Fig. 9D). Similar to the EVs, the presence of coagulation factors in BALF suggests vascular leakage and extravasation of plasma proteins into the airways that is more severe in the absence of AhR. In the protease regulation cluster, protease inhibitors such as Serpina3 and Serpina1 variants were most elevated in *Ahr^+/-^* mice, while proteins involved in coagulation and ECM remodeling, including Hrg, were more highly upregulated in BALF from *Ahr^-/-^* mice (Fig. 9E). This indicates that in response to cannabis smoke, *Ahr^+/-^* mice favor protective inhibitors, like alpha-1 antitrypsin (A1AT, Serpina1; Supp. Fig. 12A-C), a protein that helps control inflammation and limit tissue damage, whereas *Ahr^-/-^* mice have elevated BALF proteins that promote tissue breakdown and sustain inflammation.

**Figure 9.**
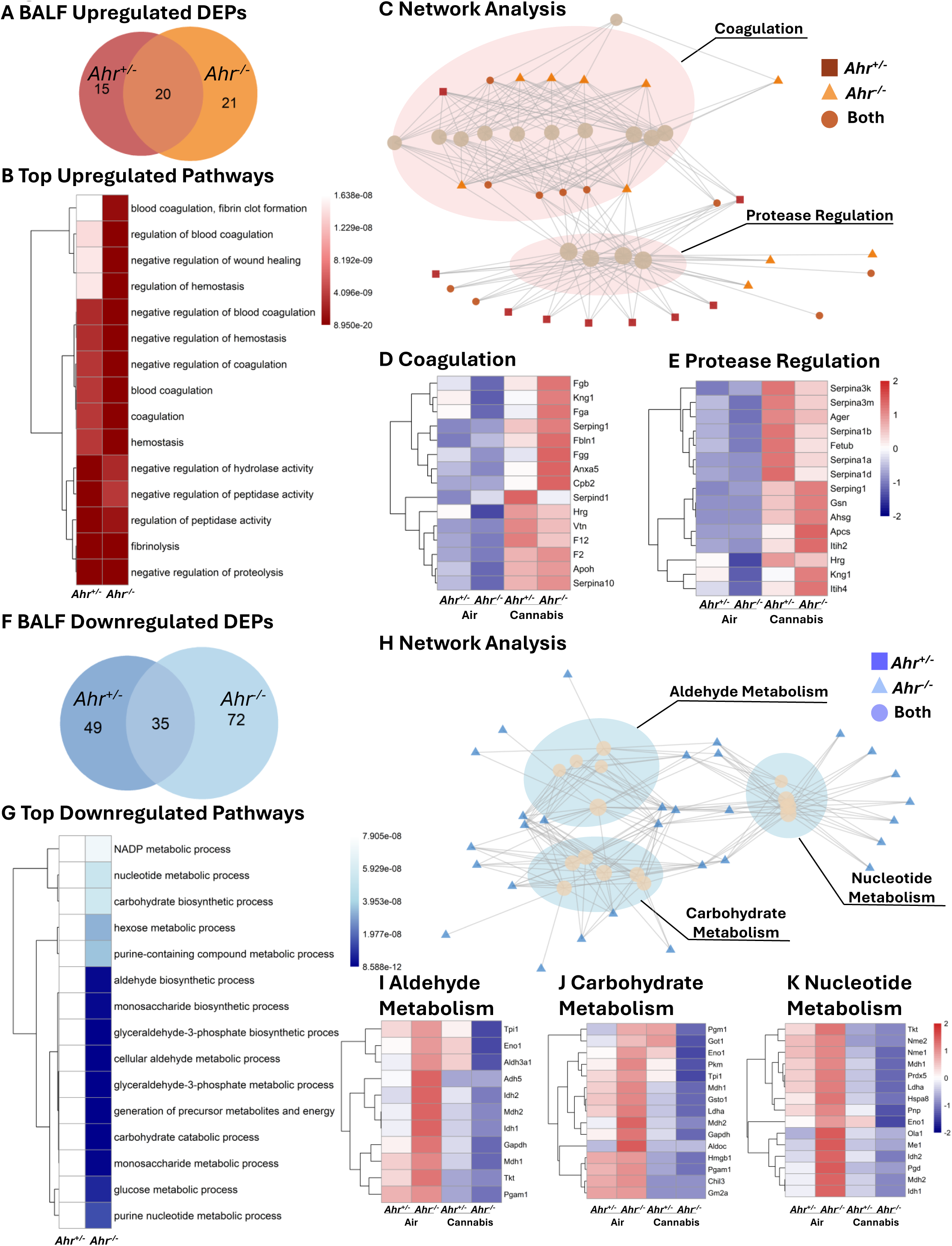
Pathway analysis of DEPs in BALF of *Ahr^+/-^* and *Ahr^-/-^* mice exposed to cannabis smoke versus air. (A) Venn diagram showing the overlap of upregulated DEPs for *Ahr^+/-^* and *Ahr^-/-^* BALF after cannabis smoke exposure versus air. (B) Top 15 upregulated pathways in *Ahr^+/-^* and *Ahr^-/-^* BALF in response to cannabis smoke exposure. (C) Network analysis of upregulated pathways. (D-E) Heatmaps of top 15 proteins in each cluster including coagulation (D) and protease regulation (E). (F) Venn diagram of downregulated DEPs for *Ahr^+/-^* and *Ahr^-/-^* BALF after cannabis smoke exposure versus air. (G) Top 15 downregulated pathways in response to cannabis smoke exposure. (H) Network analysis of downregulated pathways. (I-K) Heatmaps of top 15 proteins in the aldehyde metabolism (I), carbohydrate metabolism (J), and nucleotide metabolism (K) pathway clusters. Heatmaps represent z-scores of protein expression across conditions.

*Ahr^-/-^* mice also exhibited a greater number of BALF downregulated DEPs compared to *Ahr^+/-^* mice, and the two genotypes shared 35 downregulated DEPs (Fig. 9F, Supp. Table 24). The top 15 downregulated pathways in response to cannabis smoke exposure were observed exclusively in *Ahr^-/-^* mice (Fig. 9G, Supp. Table 25) and mostly pertained to metabolic and biosynthetic processes. Network analysis grouped these pathways into aldehyde metabolism, carbohydrate metabolism, and nucleotide metabolism (Fig. 9H, Supp. Fig. 11B). To further explore these metabolic shifts, we investigated the top 15 proteins within each category (Figs. 9I-K). In the aldehyde metabolism cluster, proteins such as Tpi1, Aldh2, Adh5, and Gapdh were highly expressed in *Ahr^-/-^* BALF at baseline but significantly decreased following cannabis smoke exposure, whereas *Ahr^+/-^* mice showed no significant changes (Fig. 9I). These proteins are involved in detoxifying reactive aldehydes and maintaining cellular redox balance, suggesting that AhR loss may impair aldehyde clearance in response to cannabis smoke. Similarly, in the carbohydrate metabolism cluster, key glycolytic and glucose metabolism proteins, including Pgm1, Eno1, and Pkm followed the same pattern of high baseline expression in *Ahr^-/-^* BALF and suppression after cannabis smoke exposure, while remaining stable in *Ahr^+/-^* mice (Fig. 9J). This suggests that AhR may play a role in maintaining carbohydrate metabolism and energy balance in the lungs following environmental exposures. In the nucleotide metabolism cluster, proteins such as Nme1, Nme2, and Pnp were also significantly reduced in *Ahr^-/-^* BALF following cannabis smoke exposure, suggesting a disruption in purine and pyrimidine metabolism (Fig. 9K). As nucleotide metabolism is critical for cellular repair, signaling, and redox homeostasis, this indicates that AhR deficiency may impair metabolic adaptation in the airways after smoke exposure. Overall, these findings suggest that AhR is required to maintain metabolic stability in BALF, particularly in aldehyde detoxification, carbohydrate metabolism, and nucleotide biosynthesis. In *Ahr^-/-^* mice, cannabis smoke exposure leads to a significant metabolic suppression, which could impair airway energy balance, oxidative stress responses, and biosynthetic processes, ultimately affecting lung homeostasis.

## Discussion

Elucidating the interplay among the various components of the respiratory system is essential for understanding the pulmonary response to environmental exposures. These exposures range from air pollution and occupational hazards to tobacco, e-cigarette aerosols, and cannabis smoke. The recent legalization of cannabis in Canada and other countries correlates with an increased rate of use (37). Currently, cannabis smoke is the third most common recreational substance after alcohol and tobacco (1). Despite its widespread use, the pulmonary effects of cannabis smoke remain largely understudied due to historical legal restrictions. Although cannabis has been suggested to have anti-inflammatory and bronchodilatory benefits due to the presence of cannabinoids such as THC (38), inhalation of smoke elicits adverse respiratory symptoms (39). One of the inherent challenges in elucidating the effects of cannabis smoke in people is concomitant tobacco use (39). In this study, we circumvent this limitation by using a preclinical model of cannabis smoke exposure that mimics human puff topography. Our study highlights significant risks associated with cannabis smoke inhalation and demonstrates that cannabis smoke induces inflammation and tissue damage, with key features including neutrophilia, vascular leakage, and tissue remodeling pathways. Moreover, these consequences of cannabis smoke are dependent on AhR, an environmental sensor that plays a pivotal role in coordinating the pulmonary response to inhaled stimuli.

A key aspect of our investigation was the analysis of EVs, which represent an emerging yet understudied component of lung biology. EVs, which includes exosomes, apoptotic bodies and microvesicles, function as critical mediators of intercellular communication by transferring molecular cargo that can regulate immune responses or cellular adaptation to environmental stressors (40). Under physiological conditions, EVs facilitate the exchange of lipids, proteins, and nucleic acids between cells. In pathological states, EVs can propagate inflammatory signals, modulate immune cell function, and contribute to the pathogenesis of lung diseases such as COPD, idiopathic pulmonary fibrosis (IPF), and acute lung injury (ALI) (41). Exposure to air pollution and cigarette smoke affects lung-derived EVs (42). However, to our knowledge, the impact of cannabis smoke on pulmonary EVs has remained unexplored, making our study the first to report that cannabis smoke significantly changes the protein cargo of lung-derived EVs. Another facet of EV biology that remains poorly characterized is the molecular mechanisms that dictate the formation and packaging of EVs. Previous work indicates that PAHs, many of which are known AhR ligands, increase EV release from hepatocytes (43). Our proteomic analysis strongly implicates a dominant role of AhR in the regulation of EV cargo, particularly proteins involved in antioxidant defense mechanisms. Notable among these were proteins related to GSH, a tripeptide thiol that can reach mM concentrations in the lung lining fluid (44). GSH is the most prevalent antioxidant in the body, playing a central role in maintaining redox balance, detoxification, and cellular defense against oxidative stress (45). GSH metabolism pathways were enriched at baseline in *Ahr^+/-^* mice but were suppressed following cannabis smoke exposure. Because the AhR interacts with nuclear factor erythroid 2-related factor 2 (NRF2), a master transcriptional regulator of oxidative stress responses and GSH biosynthesis (46), it is interesting to speculate that alterations in protein the GSH biosynthesis pathways are controlled via AhR-dependent regulation of Nrf2. However, our previous work failed to show direct regulation of Nrf2 by AhR in pulmonary cells exposed to cigarette smoke (20). Moreover, several of the proteins identified were detoxification enzymes that utilize of GSH as a cofactor (Gpx1) or are involved in GSH recycling (Idh1), rather than its production (47, 48). This may represent changes to the oxidant burden to the lung imposed by cannabis smoke that could ultimately overwhelm its antioxidant capacity. Thus, our data suggest that AhR extends its antioxidant functions to the modulation of proteins within EV cargo, influencing the extracellular redox capacity of the lung microenvironment.

In addition to proteins related to GSH metabolism and detoxification, AhR also regulated pulmonary EV proteins associated with proteostasis, a process that ensures cellular proteins are in the correct concentration, conformation, and location, and that misfolded or damaged proteins are removed. Our data shows that there is downregulation of proteins related to the 19S and 20S proteasome subunits from *Ahr^-/-^* mice, suggesting impaired protein turnover mechanisms. Soluble proteasome activity plays a crucial role in pulmonary protein homeostasis by degrading misfolded and damaged proteins, preventing their accumulation and thereby protecting against cellular stress (49, 50). Dysregulated proteasomal function has been implicated in various pulmonary diseases, including COPD and IPF, where reduced proteasome activity is associated with impaired lung function and tissue remodeling (51–53).

Tissue remodeling is a dynamic process that involves the reorganization of tissues and can either be physiological or pathological. In the lungs, remodeling involves structural changes that are often triggered by injury or inflammation, leading to diseases such as asthma, COPD, and fibrosis (54). During lung tissue remodeling, proteases play an especially important role, and thus maintaining a proper protease-antiprotease balance ensures maintenance of tissue integrity and ECM stability. Proteases such as neutrophil elastase (NE) and matrix metalloproteinases (MMPs) drive ECM degradation that facilitates tissue remodeling (55, 56), an effect that is counterbalanced by the antiproteases A1AT and tissue inhibitors of metalloproteinases (TIMPs). The function of these enzymes is necessary to preserve lung structure and prevent unwarranted tissue destruction, particularly during neutrophilic inflammation (57). Disruptions in this balance, towards favoring protease activity, cause lung damage. Exposure to cigarette smoke is a potent neutrophilic stimulus and a major cause of COPD (58). The best characterized genetic cause of COPD is A1AT deficiency (AATD) (59). AATD is caused by a mutation of the *SERPINA1* gene and affects 1/2000-1/5000 individuals (59). A1AT is produced in the liver and enters the lungs from the circulation to counteract the effects of NE released from recruited neutrophils during conditions of inflammation and infection (60). Individuals with severe AATD are predisposed to early-onset emphysema (59). We have previously shown that loss of AhR in mice leads to airspace enlargement, analogous to the emphysema component of COPD, concomitant with unrelenting neutrophilic inflammation in response to cigarette smoke exposure (21, 25, 28). Herein, we identify A1AT as a novel, AhR-regulated antiprotease, further supporting that the AhR serves as a critical protective mechanism against lung damage caused by inhaled pollutants. In *Ahr^+/-^* mice, cannabis smoke exposure induced A1AT expression, an increase that was absent in *Ahr^-/-^* mice, suggesting that AhR is required for this increase in BALF levels of A1AT. Because AhR deficiency is also associated with numerous hepatic alterations, it remains undetermined from our study whether lower levels of A1AT in *Ahr^-/-^* mice is due to decreased production by hepatocytes. Regardless, given that AATD is a risk factor for COPD (61), our findings provide the first evidence that AhR directly controls A1AT levels, offering a new perspective on how AhR protects the lung from smoke-induced damage.

Maintenance of A1AT levels is important given the heightened neutrophilic response in *Ahr^-/-^* mice exposed to cannabis smoke. Neutrophils are potent effectors of inflammation and play a crucial role in early immune responses (62). However, excessive neutrophilic infiltration can contribute to lung tissue damage through the release of serine proteases (including NE), reactive oxygen species, and inflammatory cytokines (63). Elevated neutrophils were observed in the blood and lung parenchyma after cannabis smoke exposure, with *Ahr^-/-^* mice exhibiting significantly stronger neutrophilic response. It is interesting that there was no increase in neutrophils in the airways, indicating a compartmentalized inflammatory response to cannabis smoke. This pattern differs from tobacco smoke exposure, which is associated with an influx of neutrophils into the airways (28) and suggests that cannabis smoke primarily affects the interstitium and vasculature of the lung. This pattern may result from dysregulated neutrophil trafficking, potentially driven by alterations in adhesion molecule expression, rather than changes in chemotactic cytokines. We draw this conclusion because despite analysis of dozens of cytokines by multiplex array, there were few significant differences between cannabis smoke-exposed *Ahr^+/-^* and *Ahr^-/-^* mice. Furthermore, cytokines such as IL-6 were elevated in BALF from both *Ahr^+/-^* and *Ahr^-/-^* mice, indicating a localized inflammatory response. However, cannabis smoke significantly downregulated key adhesion molecules involved in neutrophil diapedesis, including Pecam1, Icam2, and Esam. These proteins are essential for neutrophil tethering to endothelial cells and subsequent transmigration through endothelial and epithelial layers into the airspaces (64). Given that cannabis smoke reduces the expression of these proteins, particularly in *Ahr^+/-^* mice, this impairment can explain the absence of neutrophils in the airways despite their accumulation in the vasculature and interstitium. This highlights a unique inflammatory signature associated with cannabis smoke exposure and underscores the critical role of AhR in regulating neutrophil recruitment to distinct lung compartments.

An emerging theme throughout this work is that AhR may primarily protect the lung by preserving epithelial barrier and structural integrity. Disruption of this barrier can predispose individuals to a range of health complications, including increased susceptibility to infection (65). When epithelial integrity is compromised, pathogens can penetrate deeper into lung tissue and even enter the circulation, increasing the risk of systemic infection (66). In parallel, barrier disruption amplifies inflammatory signaling and immune cell infiltration, leading to excessive inflammation, pulmonary edema, and impaired gas exchange (67). It is therefore notable that in addition to controlling anti-protease activity, our data revealed that AhR deficiency leads to a disruption in epithelial barrier integrity following cannabis smoke exposure, evidenced by significantly higher epithelial cell sloughing in the BALF from *Ahr^-/-^* mice. Mechanistically, this observation is supported by lower expression of adhesion molecules and cytoskeletal regulators in lung tissue from *Ahr^-/-^* mice, pointing to impaired cell–cell contact and cytoskeletal stability as key consequences of AhR deficiency. Previous studies have implicated AhR in maintaining endothelial integrity (66) and our findings extend this role to the lung epithelium, where AhR appears to be essential for maintaining structural cohesion under conditions of environmental stress. The increased vascular permeability observed in *Ahr^-/-^* mice, inferred from elevated coagulation factors in BALF and EVs, further suggests that AhR contributes to pulmonary endothelial and epithelial homeostasis. These effects may be mediated by AhR-dependent transcriptional regulation of genes involved in tight junction assembly, adherens junction stability, or actin cytoskeletal organization; several of which were dysregulated in our dataset.

Despite the valuable insights provided by this study, several limitations warrant consideration. The use of bulk proteomics limits resolution at the level of individual cell types, which is a significant constraint given the cellular heterogeneity of lung tissue. Incorporating single cell or spatial proteomic approaches in future studies could help define the specific cellular origins of observed proteomic changes. Moreover, this study focused on acute cannabis smoke exposure, capturing early molecular responses but not the long-term or pathological consequences of chronic exposure. Chronic inhalation better reflects typical patterns of cannabis use in humans and may result in cumulative effects on lung structure, function, and disease progression. Although our findings implicate AhR in the regulation of pathways involved in barrier integrity, proteostasis, and antioxidant defense, we did not directly investigate the molecular drivers of these changes. Follow-up studies are needed to dissect the specific signaling pathways and gene regulatory networks responsible for AhR-mediated protection. Finally, our exposure model used a single high THC cannabis cultivar; however, cannabis is a chemically diverse plant with thousands of strains varying in cannabinoid and terpene content. As a result, the findings may not be generalizable across cannabis products with different chemical profiles or modes of delivery, such as vaping or oral ingestion. Broader investigation into the pulmonary effects of other cannabis formulations will be essential to inform evidence-based health risk assessments and regulatory policies.

Despite its notoriety as a driver of toxic responses to man-made chemicals, the AhR has emerged as a key player in the homeostatic regulation of organ function. Overall, our study provides new, critical insights into the AhR-dependent modulation of the pulmonary effects of cannabis smoke, which remain poorly understood despite worldwide cannabis use. While cannabis is widely perceived as safe, or even therapeutic, our data highlight distinct and significant risks associated with its use, including inflammation and damage to the epithelial layer. Given the increasing prevalence of cannabis use, including the preference for smoking, future studies should investigate the long-term consequences of cannabis exposure on respiratory health. These findings provide mechanistic insights into the biological consequences of cannabis smoke exposure and emphasize the importance of AhR signaling in mitigating pulmonary damage. As cannabis use continues to rise globally, understanding the molecular drivers and genetic factors that influence susceptibility to its pulmonary effects will be critical for guiding evidence-based public health recommendations.

## Data availability

All data are available within this article, the supplemental file, online at https://figshare.com/s/b998473f7720cc617317 and from the corresponding author upon reasonable request.

## Author Contributions

ETW and CJB were responsible for conceptualization and project administration. Exposures and sample generation were performed by ETW. Flow cytometry was performed by ETW and RG. EV extraction was performed by ETW, AB, and TT. Data analysis was conducted by ETW and NSH. The first draft of the manuscript was written by ETW. All authors reviewed and commented on previous versions of the manuscript and approved the final version.

## Conflict of Interest Statement

The authors declare no conflicts of interest.

## Acknowledgments

This project was supported by funding acquired by CJB from the Canadian Institutes of Health Research Project Grant 162273. CJB was supported by the Fonds de Recherche du Québec -Santé (FRQS). The authors would also like to acknowledge the Proteomics and Molecular Analysis Platform of the Research Institute of the McGill University Health Centre for their technical support.

## Notes

https://figshare.com/s/b998473f7720cc617317

### Competing Interest Statement

The authors have declared no competing interest.

