## Supplemental Figures for "The AhR is a Critical Regulator of the Pulmonary Response to Cannabis Smoke"

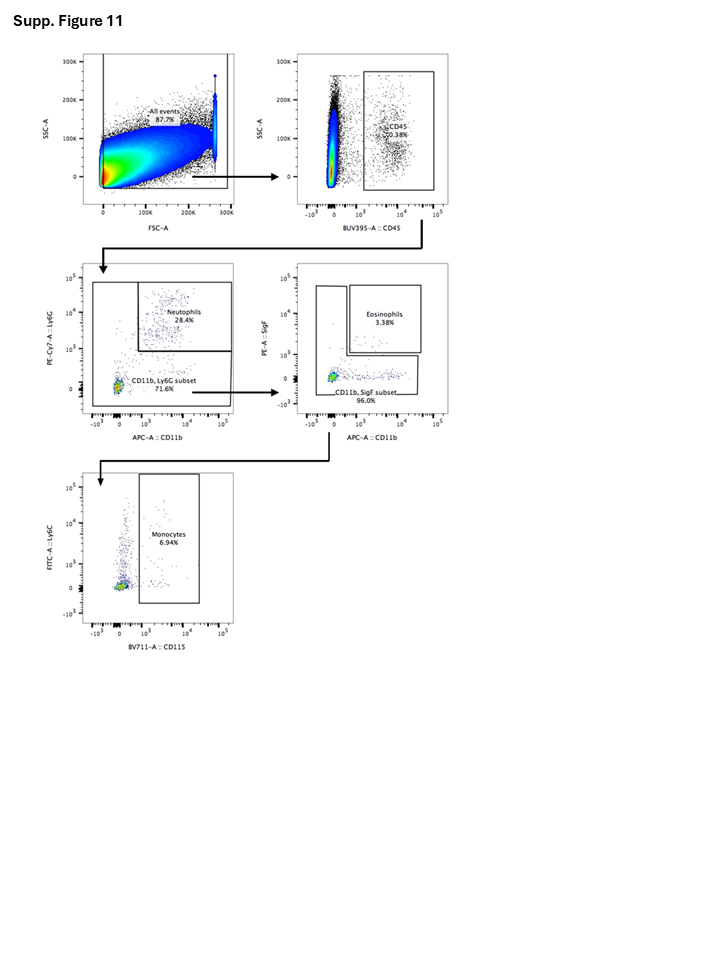

**Supp. Figure 1 Gating Strategy for Innate Immune Cells in the Blood**
Total events were selected using forward scatter (FSC-A) and side scatter (SSC-A) parameters. CD45⁺ leukocytes were gated to exclude non-immune cells. Among CD45⁺ cells, neutrophils were identified as CD11b⁺Ly6G⁺ cells, while the CD11b⁺Ly6G⁻ subset was further analyzed for eosinophils and monocytes. Eosinophils were gated based on SiglecF (SigF) expression, and monocytes were identified as CD115⁺Ly6C⁺ cells.

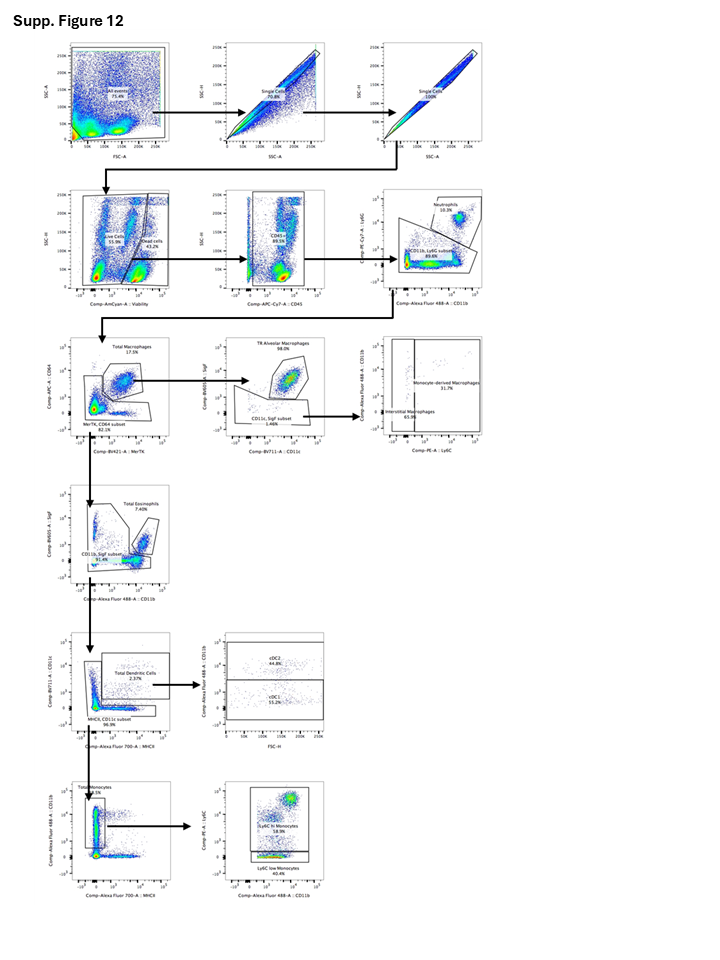

**Supp. Figure 2 Gating Strategy for Innate Immune Cells of the Lung Tissue**
Total events were selected using forward scatter (FSC-A) and side scatter (SSC-A) parameters to remove debris. Single cells were identified using SSC-A vs. SSC-H and FSC-A vs. FSC-H plots. Viable CD45⁺ immune cells were gated to exclude dead cells and non-immune populations. Among CD45⁺ cells, neutrophils were identified as CD11b⁺Ly6G⁺, while the negative subset was further analyzed for macrophages, eosinophils, dendritic cells, and monocytes. Macrophages were classified into total macrophages (MerTK⁺CD64⁺), tissue-resident (TR) alveolar macrophages (CD11c⁺SiglecF⁺), monocyte-derived macrophages (CD11c⁻SiglecF⁻Ly6C⁺), and interstitial macrophages (CD11c⁻SiglecF⁻Ly6C^-^). Eosinophils were identified as MerTK⁻CD64⁻SigF⁺CD11b⁺. DCs were classified as MHCII⁺CD11c⁺, further subdivided into type 1 conventional DCs (cDC1; CD11b⁺) and type 2 conventional DCs (cDC1; CD11b⁻). Monocytes were subdivided into Ly6C^hi^ inflammatory monocytes and Ly6C^low^ patrolling monocytes.

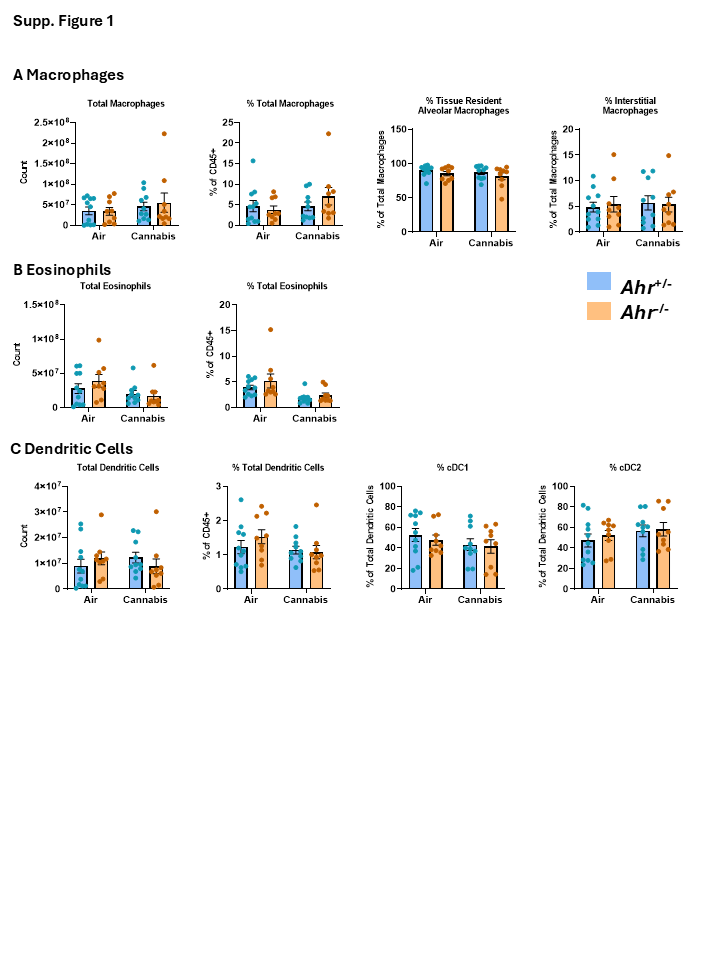

**Supp. Figure 3 Quantification of Innate Immune Cell Populations in the Lung**

(A) Macrophage populations, including total macrophage counts, percentage of total macrophages among CD45⁺ cells, percentage of total macrophages that are tissue-resident alveolar macrophages, and percentage of total macrophages that are interstitial macrophages. (B) Eosinophil populations, including total eosinophil counts and percentage of CD45⁺ cells. (C) Dendritic cell populations, including total dendritic cell counts, percentage of CD45⁺ cells, and percentage of total dendritic cells that are cDC1 or cDC2. Statistical significance is shown as *p < 0.05, **p < 0.01, ***p < 0.001, and ****p < 0.0001.

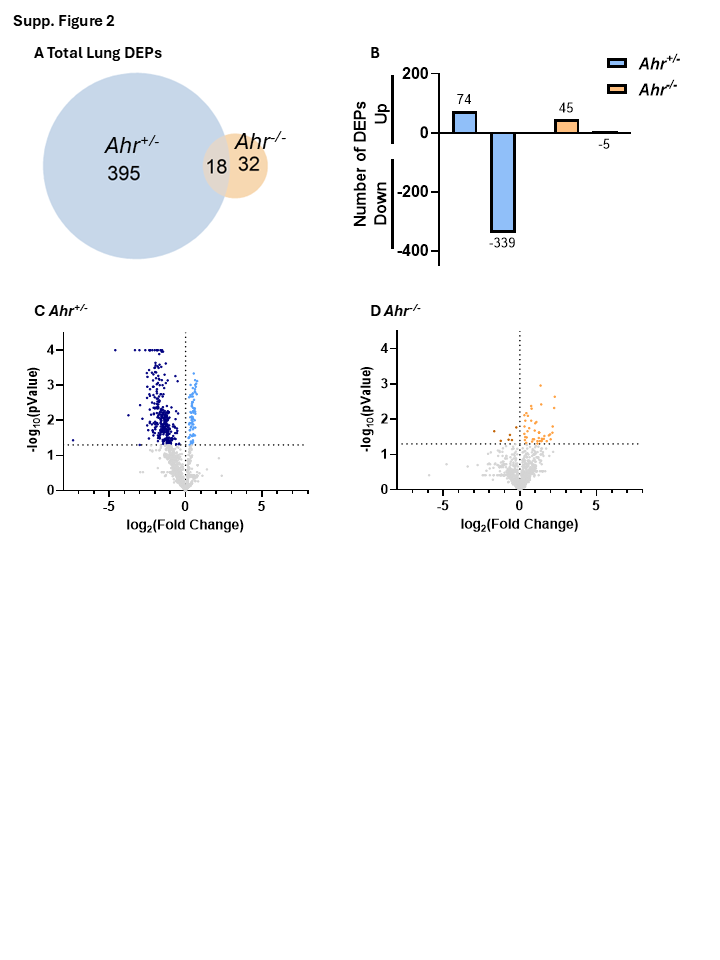

**Supp. Figure 4 Lung Tissue DEPs in *Ahr^+/-^* and *Ahr^-/-^* Mice Exposed to Cannabis Smoke vs Air**

(A) Venn diagram displaying the total number of DEPs in *Ahr^+/-^* and *Ahr^-/-^* mice exposed to cannabis smoke compared to air-exposed controls. (B) Bar graph quantifying the number of DEPs that are upregulated or downregulated in each genotype. (C, D) Volcano plots illustrating the distribution of DEPs in (C) *Ahr^+/-^* and (D) *Ahr^-/-^* mice. The horizontal dotted line indicates the statistical significance threshold (*p*<0.05).

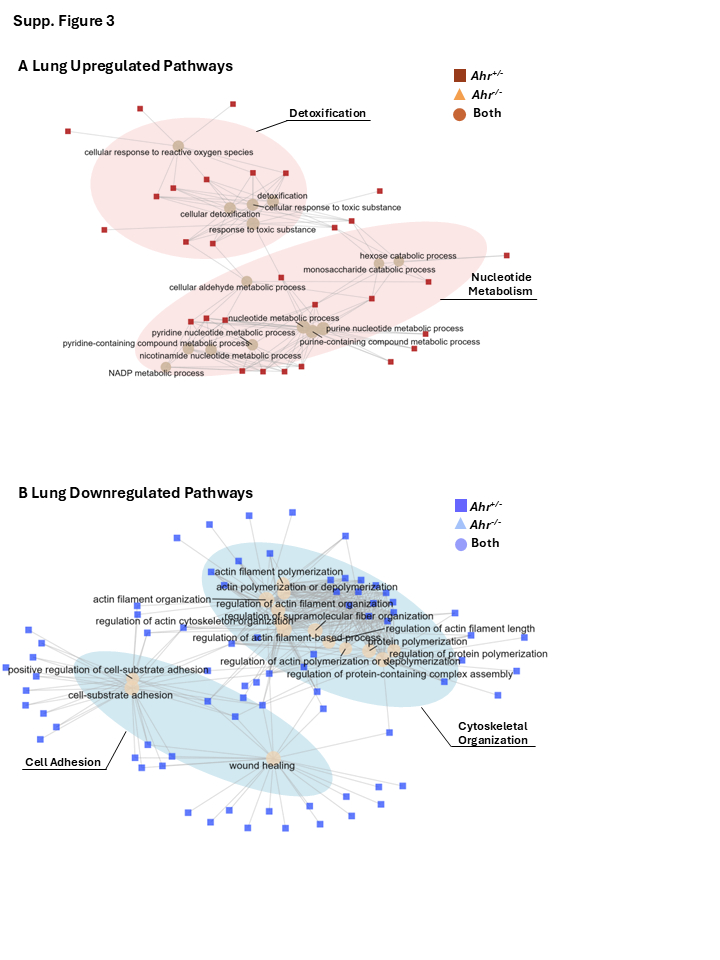

**Supp. Figure 5 Labelled Network Analysis of Top 15 Upregulated and Downregulated Pathways in Lung Tissue**

(A) Network analysis of the top 15 upregulated pathways in lung tissue following cannabis smoke exposure. (B) Network analysis of the top 15 downregulated pathways in lung tissue following cannabis smoke exposure. Beige pathway nodes are labeled with the GO term they represent, while the protein nodes are color-coded based on the genotype in which they are differentially expressed.

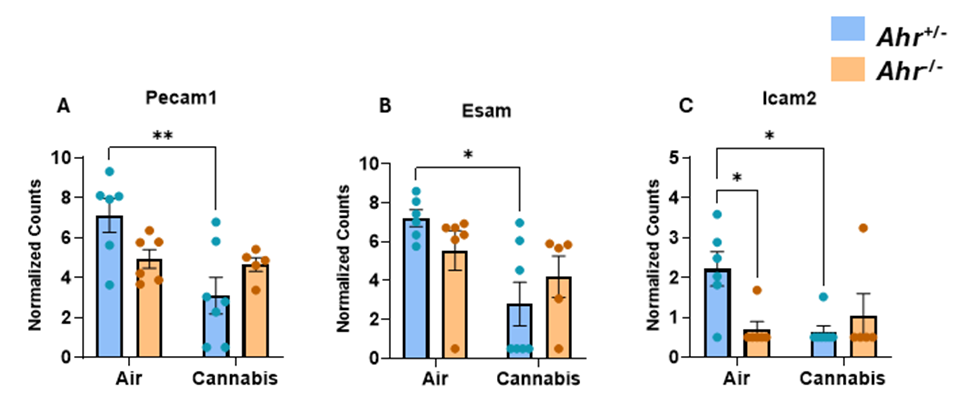

**Supp. Figure 6 Levels Adhesion Molecules involved in Diapedesis in Lung Tissue of *Ahr^+/-^* and *Ahr^-/-^* Mice Exposed to Air or Cannabis Smoke.**

Normalized counts of (A) Pecam1, (B) Esam, and (C) Icam2 in lung tissue. Data are presented as mean ± SEM, with individual data points shown. Statistical significance is denoted by *p < 0.05, and **p < 0.01.

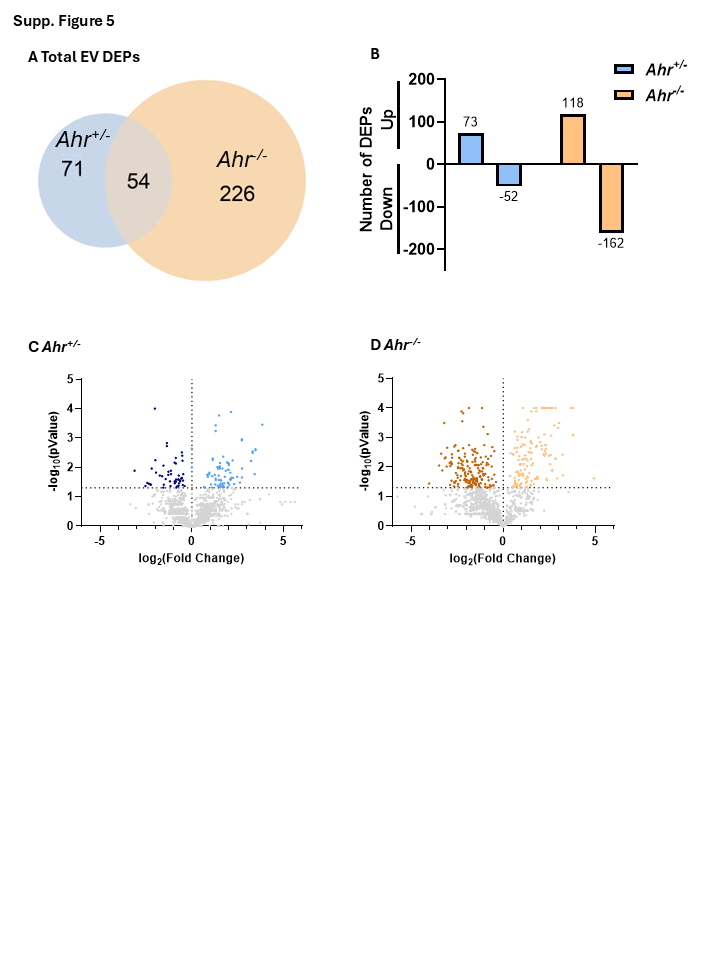

**Supp. Figure 7 EV DEPs in *Ahr^+/-^* and *Ahr^-/-^* Mice Exposed to Cannabis Smoke vs Air**

(A) Venn diagram displaying the total number of DEPs in *Ahr^+/-^* and *Ahr^-/-^* mice exposed to cannabis smoke compared to air-exposed controls. (B) Bar graph quantifying the number of DEPs that are upregulated or downregulated in each genotype. (C, D) Volcano plots illustrating the distribution of DEPs in (C) *Ahr^+/-^* and (D) *Ahr^-/-^* mice. The horizontal dotted line indicates the statistical significance threshold (*p*<0.05).

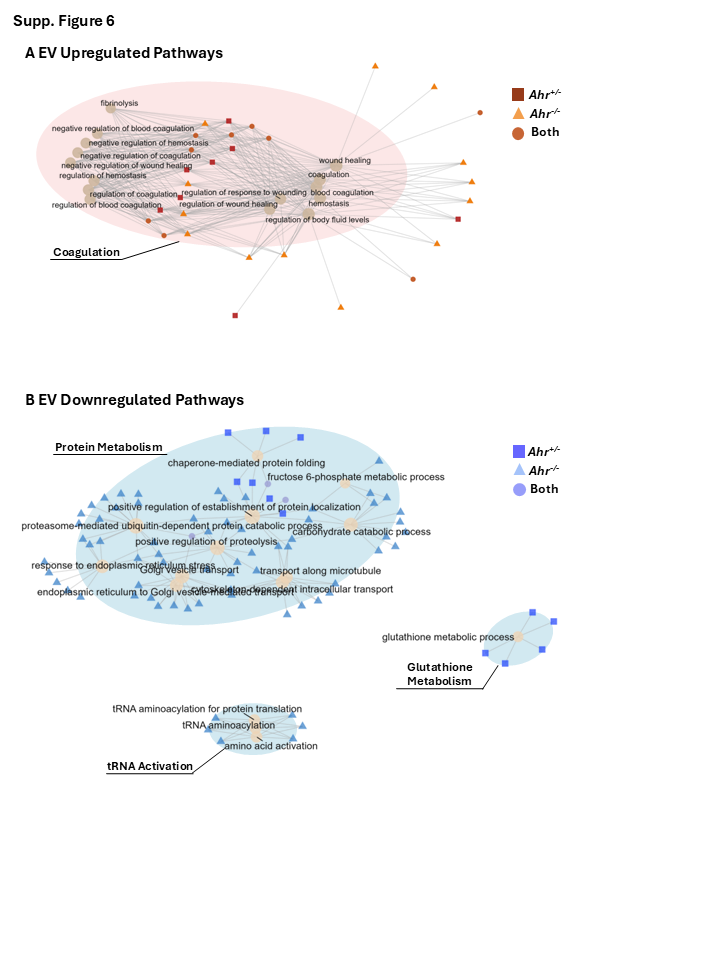

**Supp. Figure 8 Labelled Network Analysis of Top 15 Upregulated and Downregulated Pathways in EVs.**

(A) Network analysis of the top 15 upregulated pathways in EVs following cannabis smoke exposure. (B) Network analysis of the top 15 downregulated pathways in EVs following cannabis smoke exposure. Beige pathway nodes are labeled with the GO term they represent, while the protein nodes are color-coded based on the genotype in which they are differentially expressed.

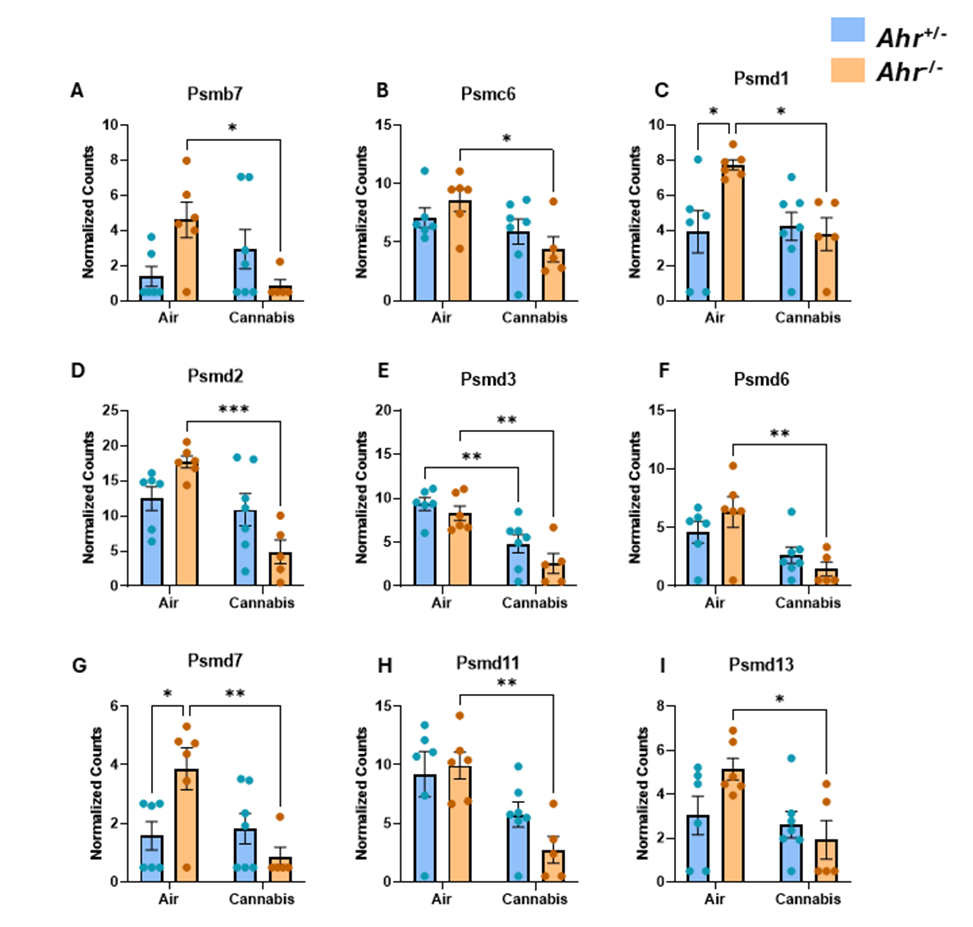

**Supp. Figure 9 Proteasome Subunits in EVs From *Ahr^+/-^* and *Ahr^-/-^* Mice Exposed to Either Air or Cannabis Smoke.**

Normalized counts for (A) Psmb7, (B) Psmc6, (C) Psmd1, (D) Psmd2, (E) Psmd3, (F) Psmd6, (G) Psmd7, (H) Psmd11, and (I) Psmd13. Data are presented as mean ± SEM, with individual data points shown. Statistical significance is denoted by *p < 0.05, **p < 0.01, and ***p < 0.001.

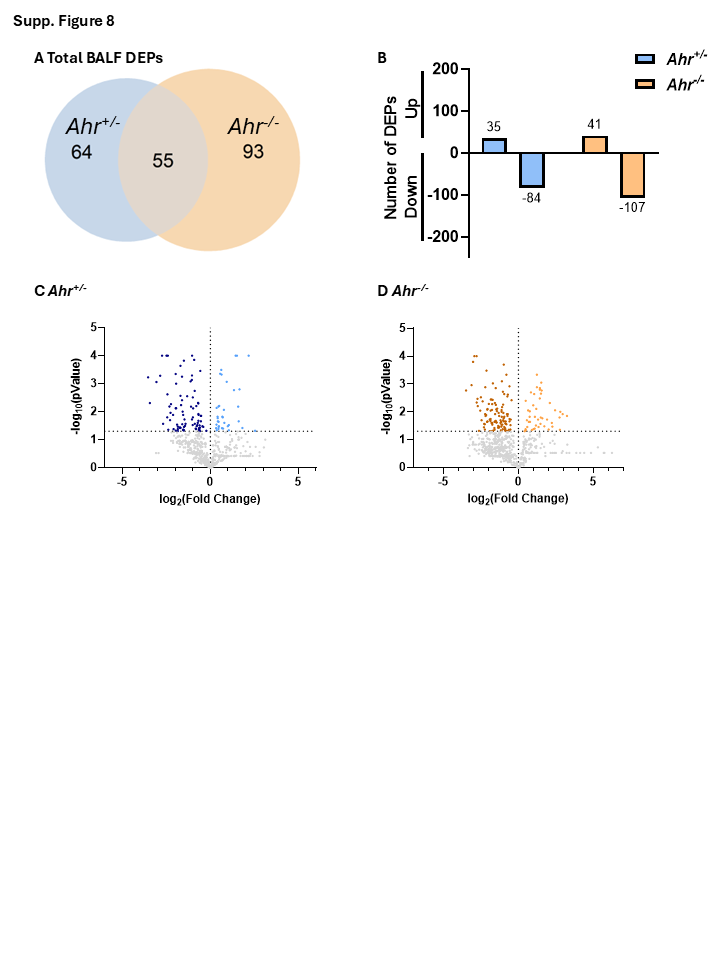

**Supp. Figure 10 BALF DEPs in *Ahr^+/-^* and *Ahr^-/-^* Mice Exposed to Cannabis Smoke vs Air**

(A) Venn diagram displaying the total number of DEPs in *Ahr^+/-^* and *Ahr^-/-^* mice exposed to cannabis smoke compared to air-exposed controls. (B) Bar graph quantifying the number of DEPs that are upregulated or downregulated in each genotype. (C, D) Volcano plots illustrating the distribution of DEPs in (C) *Ahr^+/-^* and (D) *Ahr^-/-^* mice. The horizontal dotted line indicates the statistical significance threshold (*p*<0.05).

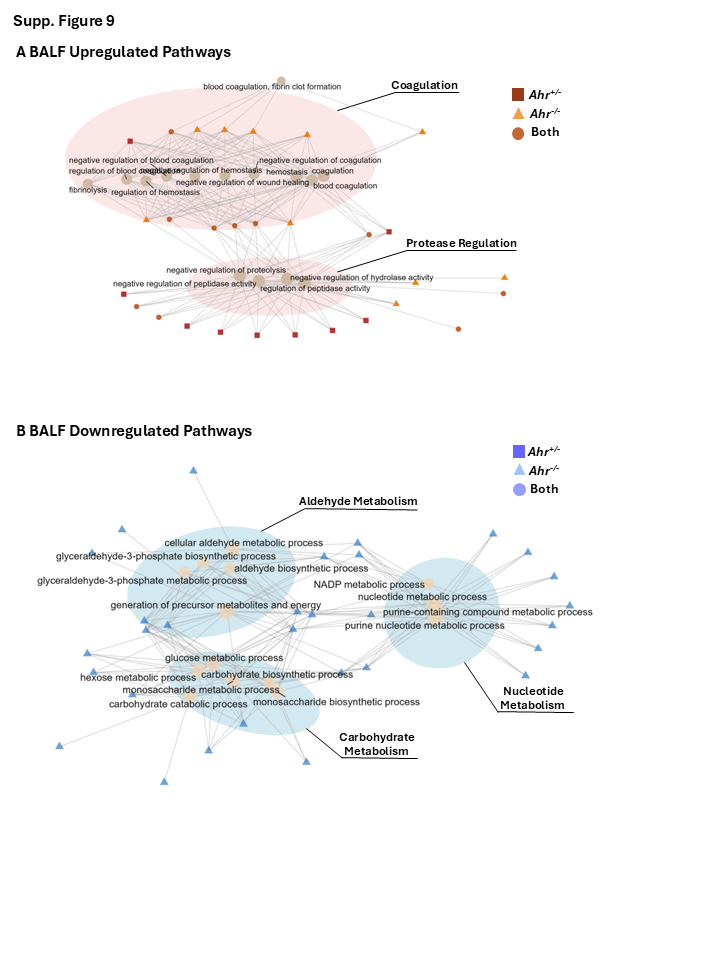
 **Supp. Figure 11 Labelled Network Analysis of Top 15 Upregulated and Downregulated Pathways in BALF.**

(A) Network analysis of the top 15 upregulated pathways in BALF following cannabis smoke exposure. (B) Network analysis of the top 15 downregulated pathways in EVs following cannabis smoke exposure. Beige pathway nodes are labeled with the GO term they represent, while the protein nodes are color-coded based on the genotype in which they are differentially expressed.

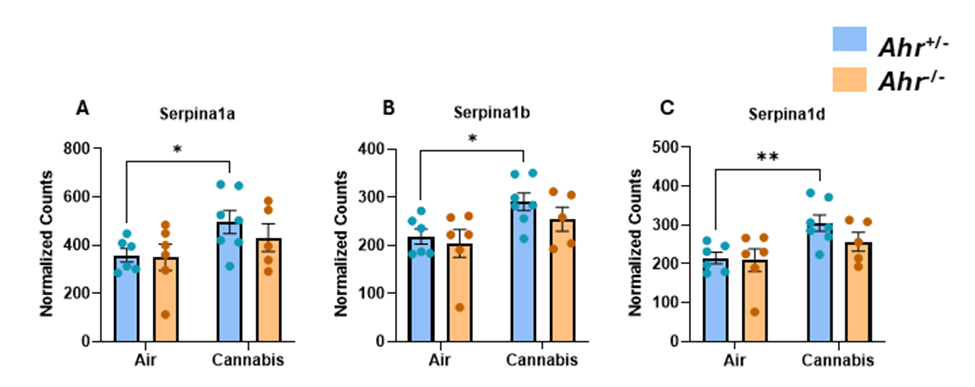

**Supp. Figure 12 Levels of A1AT Variants in BALF of *Ahr^+/-^* and *Ahr^-/-^* Mice Exposed to Air or Cannabis Smoke.**

Normalized counts of (A) Serpina1a, (B) Serpina1b, and (C) Serpina1d in BALF. Data are presented as mean ± SEM, with individual data points shown. Statistical significance is denoted by *p < 0.05, and **p < 0.01.
